# Molecular organization of the *Chlorella sorokiniana* pyrenoid

**DOI:** 10.64898/2026.07.17.739135

**Authors:** Mihris Ibnu Saleem Naduthodi, James Barrett, Jessica Pritchard, Philippe Van der Stappen, Manon Demulder, Tommaso Garfagnini, Adam Dowle, Maria Hondele, Benjamin D. Engel, Alistair McCormick, Luke C. M. Mackinder

## Abstract

To overcome the enzymatic limitations of Rubisco, algae operate CO2-concentrating mechanisms (CCMs) that deliver concentrated CO2 to Rubisco tightly packaged in a specialized microcompartment called a pyrenoid. Pyrenoids are globally important biomolecular condensates, but their convergent evolution means that their molecular composition and emergent architecture cannot be inferred across clades. Here we characterize the pyrenoid of the Trebouxiophyceae alga, Chlorella sorokiniana. Using cryo-electron tomography, we provide an architectural overview of the pyrenoid and visualize pyrenoid-specific protein complexes. Quantitative proteomics and Rubisco co-immunoprecipitation followed by mass spectrometry demonstrate that inorganic carbon delivery machinery is conserved across green algae but the pyrenoid structural components are not. In vitro reconstitution supports the role of two previously undescribed proteins, one in assembly of pyrenoid traversing thylakoids (putative matrix thylakoid tether; PMTT) and another in starch tethering to the Rubisco matrix (putative matrix starch tether; PMST). In Nicotiana benthamiana, PMTT localized to the thylakoid stromal lamellae and PMST to chloroplast starch granules. Our findings provide insights into the molecular logic of pyrenoid assembly; how proteins mediate condensate-membrane and condensate-starch interactions; and expands the pyrenoid plant engineering toolkit, setting the stage for engineering a Chlorella pyrenoid into plants.

## Introduction

Atmospheric carbon dioxide (CO_2_), the primary source of carbon for photoautotrophic organisms, is fixed into organic carbon by the protein ribulose-1,5-bisphosphate carboxylase/oxygenase (Rubisco) (Bar-On and Milo, 2019). Rubisco is catalytically slow (Bar-Even et al., 2011), and does not efficiently differentiate between CO_2_ and O_2_ as the substrate (Tcherkez, Farquhar and Andrews, 2006). The oxygenation reaction of Rubisco accumulates toxic 2-Phosphoglycolate, which is reassimilated with high energy cost. These shortcomings of Rubisco have resulted in plants producing Rubisco in large quantities to meet required carboxylation rates for competitive growth (Raven, 2013). To overcome Rubisco’s limitations during geological periods of low atmospheric CO_2_ photosynthetic organisms such as eukaryotic algae, some plants and cyanobacteria evolved CO_2_-concentrating mechanisms (CCMs) that accumulate CO_2_ in the vicinity of Rubisco to increase the CO_2_:O_2_ ratio and favour the carboxylation reaction of the enzyme (Barrett et al., 2026). A large portion of CCMs in aquatic phototrophs are mediated through an organelle called the pyrenoid, a biomolecular condensate in the chloroplast of eukaryotic algae (Freeman Rosenzweig et al., 2017; Barrett, Girr and Mackinder, 2021). It is estimated that around two-thirds of oceanic carbon fixation is channelled through the pyrenoid, indicating its importance in biogeochemical cycles (Mackinder et al., 2016). The efficiency of pyrenoids has resulted in considerable interest in engineering a pyrenoid-based CCM (pCCM) into C3 crops, such as rice, wheat and soy, which lack CCMs and where a pCCM is proposed to enhance yields and improve water and nitrogen use efficiencies (Fei et al., 2022; Long, Marshall-Colon and Zhu, 2015; Long et al., 2025).

The best studied model of the pCCM is that of green microalga *Chlamydomonas reinhardtii*. pCCM components can be split into three categories: uptake and transport of inorganic carbon (Ci; predominantly in the form of CO_2_ or bicarbonate), regulatory components, and pyrenoid structural components. Ci transport from the extracellular environment to the pyrenoid in *Chlamydomonas* is mediated by multiple proteins. HLA3 imports bicarbonate from outside the cell into the cytosol, after which it is transported into the chloroplast by LCIA (Gao et al., 2015; Yamano et al., 2015). Within the chloroplast, bestrophin-like proteins (BSTs) enriched at the pyrenoid periphery channel bicarbonate into pyrenoid traversing thylakoids (PTTs) where the carbonic anhydrase, CAH3, converts bicarbonate to CO_2_, which then diffuses into the pyrenoid matrix for fixation by Rubisco (Mukherjee et al., 2019; Sinetova et al., 2012). Stromal localized carbonic anhydrases, LCIB and LCIC, play an essential role in the CCM by generating bicarbonate from CO_2_ that passively enters into the chloroplast or has leaked from the pyrenoid (Yamano et al., 2010). In addition to the transporters and carbonic anhydrases, CAS a calcium binding protein localized to the PTTs, and CIA5 a nuclear localized transcriptional regulator, regulate the pCCM in *Chlamydomonas* (Wang et al., 2016; Wang et al., 2015). Following Ci delivery, multiple components are required for the structural integrity of the pyrenoid; these include EPYC1 for Rubisco condensation to form the matrix (Mackinder et al., 2016), MITH1, SAGA1 and BST4 for PTT formation (Hennacy et al., 2024; Garde, Wu and Jonikas, 2026), and SAGA1, SAGA2, and EPOS1 for starch sheath formation (Crans et al., 2026; Adler et al., 2026). Additionally, multiple other pyrenoid-associated proteins have been identified, with yet to be verified functions (Lau et al., 2023; Meyer et al., 2020; Mackinder et al., 2017; Wang et al., 2023)

The recent progress in understanding *Chlamydomonas* pCCM function has guided the heterologous expression of components in plants, resulting in the formation of a proto-pyrenoid matrix (Atkinson et al., 2020) and assembly of additional structural elements such as starch attachment to the proto-pyrenoid and PTTs (Hennacy et al., 2024; Atkinson et al., 2024). However, several challenges remain in engineering a functional pyrenoid in crops. In *Chlamydomonas*, the linker protein EPYC1, which drives pyrenoid formation through biomolecular condensation of Rubisco, binds specifically to the *Chlamydomonas* small subunit of Rubisco (CrRbcS) (Mackinder et al., 2016; He et al., 2020) . However, due to RbcS sequence divergence, EPYC1 is not compatible with the RbcS of C3 plants (Barrett et al., 2024). Thus, plant engineering approaches require substitution of the endogenous RbcS with CrRbcS to enable proto-pyrenoid assembly (Atkinson et al., 2020). In addition, the *Chlamydomonas* pyrenoid exhibits considerable structural complexity, including multiple PTTs that undergo large structural rearrangements and contain poorly understood internal minitubules, and a surrounding starch sheath composed of multiple plates (Engel et al., 2015). There has recently been significant progress in understanding both PTT and starch-sheath formation (Hennacy et al., 2024; Crans et al., 2026; Adler et al., 2026; Garde, Wu and Jonikas, 2026; Wu et al., 2026), however, there are still many unknowns that complicate efforts to reconstitute these structures in plants. Consequently, characterizing pyrenoids from other microalgal lineages with simpler architectures and potentially fewer or less specific components could accelerate progress toward engineering pyrenoids into crops (Adler et al., 2022; Pritchard et al., 2026; Barrett et al., 2026).

The fast-growing green microalga, *Chlorella sorokiniana* (*Chlorella* from here on), has a morphologically distinct and simpler pyrenoid, which we previously showed contains a linker protein (CsLinker) that can directly induce the condensation of a broad range of Rubiscos (Barrett et al., 2024). Here, we set out to characterize the CCM and pyrenoid of *Chlorella* to improve our fundamental understanding of pyrenoid assembly in a different algal clade and to lay the foundation for *Chlorella* pyrenoid engineering in plants. We explore *Chlorella* pyrenoid architecture via cryo-electron tomography (cryo-ET) and decipher the pCCM via proteomics to identify three novel pyrenoid proteins. We perform detailed in vitro and in vivo characterization of these newly identified core pyrenoid proteins, revealing their potential roles in *Chlorella* pyrenoid organization. Expression of these proteins *in planta* resulted in functional localization, setting the stage for developing a *Chlorella* pyrenoid in plants.

## Results

### *Chlorella* has a low CO_2_-inducible pyrenoid-based CCM

*Chlorella* belongs to the green algal class Trebouxiophyceae, which diverged around 850 million years ago (Ma) from the Chlorophyceae, the class that includes *Chlamydomonas* (Del Cortona et al., 2020). Considering this ancient divergence, we set out to characterize the pCCM of *Chlorella* and compare it to the well-studied *Chlamydomonas* pCCM.

We initially confirmed that *Chlorella*, like *Chlamydomonas*, has an inducible CCM which we assessed by cellular affinity to inorganic carbon (Ci) of cells grown under deplete or replete Ci levels. We measured the oxygen evolution rates for *Chlorella* and, as a control, *Chlamydomonas* cultures adapted to two depleted Ci levels (very low CO_2_, VLC, ∼100 ppm; and low CO_2_, LC, ∼400 ppm), and one replete Ci level (high CO_2_, HC, ∼30,000 ppm). VLC and LC-grown *Chlorella* showed significantly higher affinity for inorganic carbon compared to cultures adapted to HC (Fig. 1a, b; Extended Data Fig. 1). To assess if CCM-induction was accompanied by cellular structural changes, we performed transmission electron microscopy (TEM) analysis of *Chlorella* cells adapted to HC, LC and VLC conditions. A single distinct pyrenoid with a singular PTT and a starch-sheath encapsulating the matrix was clearly observed in LC and VLC adapted cells. This contrasts to HC-adapted cells, which had starch distributed throughout the chloroplast and typically lacked an intact pyrenoid confirming that the pyrenoid is induced in *Chlorella* at LC and VLC conditions (Fig. 1c; Extended Data Fig. 2a,b). Although in a small proportion of sections at HC, we observed a reduced pyrenoid matrix that lacked a starch-sheath and contained a PTT (Extended Data Fig. 2b,c,d).

**Figure 1.**
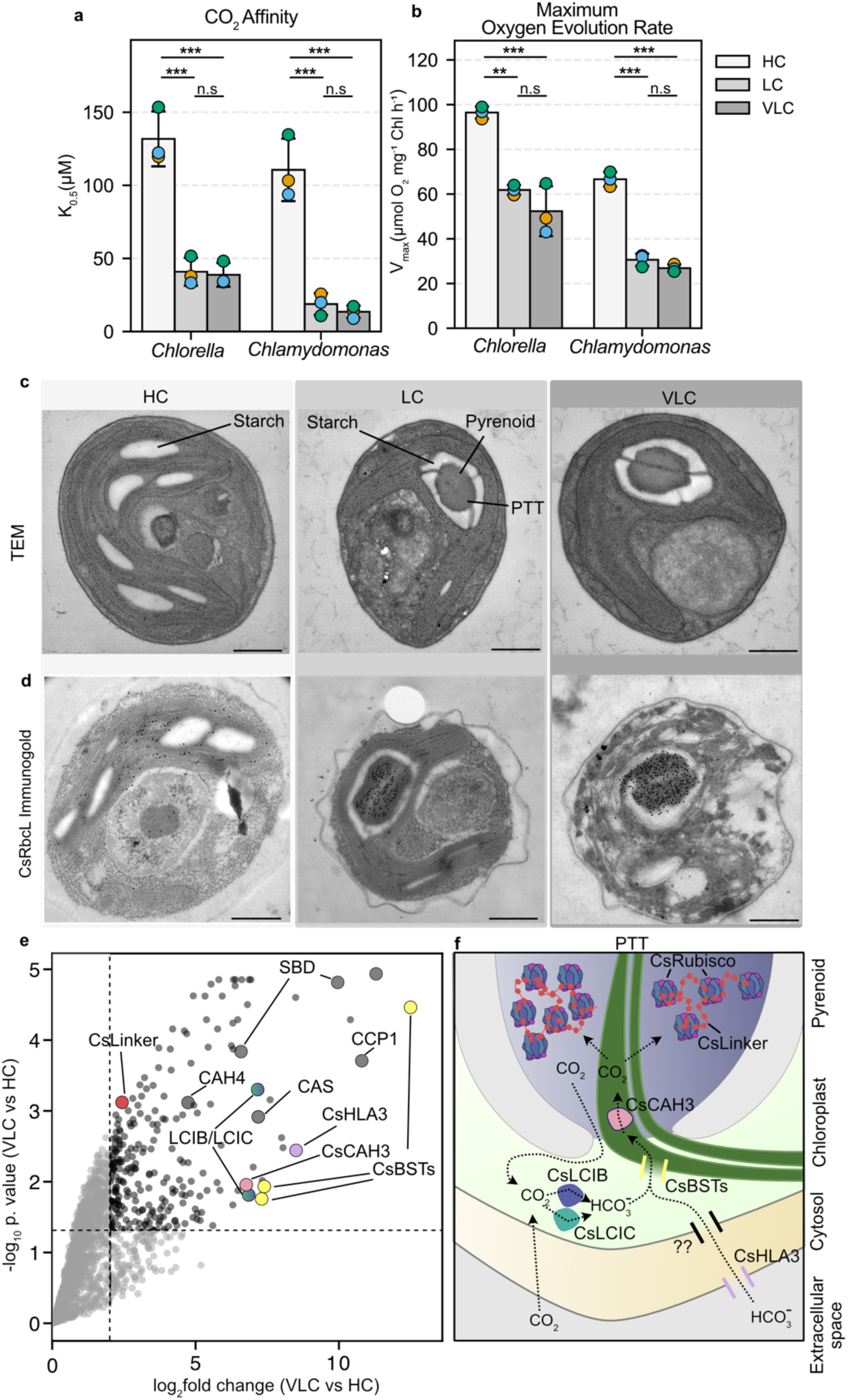
*Chlorella* has an inducible pyrenoid-based CCM. a) K_0.5_ and b) V_max_ values for *Chlorella* and *Chlamydomonas* cultures adapted to high CO_2_ (HC) and low CO_2_ (LC) conditions. Data are presented as mean ± SD (n = 3 independent biological replicates). Statistical significance between conditions within each strain was determined using a one-way ANOVA followed by Tukey’s honestly significant difference (HSD) post-hoc test. Adjusted p-values are indicated as follows: n.s (not significant, p ≥0.05), * (p < 0.05), ** (p < 0.01), and *** (p < 0.001). c) Transmission electron microscopy (TEM) of resin-embedded *Chlorella* cells grown in HC, LC and VLC. Scale bars, 500 nm. d) TEM immunogold labelling of Rubisco in *Chlorella* cells adapted to HC, LC and VLC. Gold particles are enlarged for visualization and example raw images are shown in Extended Data Fig. 3. Scale bar, 500 nm. e) Comparative proteomics of *Chlorella* cells adapted to very low CO_2_ (VLC) and high CO_2_ HC. CsLinker, starch binding domain (SBD) containing proteins and proteins with sequence conservation to characterized inorganic carbon delivery components in *Chlamydomonas* are highlighted (see Supplementary Tables 1 and 2). f) Proposed schematic of inorganic carbon transport machinery found in *Chlorella* along with known pyrenoid components Rubisco and CsLinker. Based on comparative proteomics data.

Immunogold labelling of these cells with an antibody raised to a Rubisco large subunit (RbcL) peptide showed the localization of Rubisco within the pyrenoid matrix at LC and VLC, while a diffuse distribution was observed in the chloroplast at HC (Fig. 1d, Extended Data Fig. 3). Our TEM data indicates that pyrenoid formation is dynamic and responsive to CO_2_ levels in *Chlorella*, and accompanies an increase in Ci affinity, consistent with the induction of the pCCM in *Chlamydomonas* (Moroney and Ynalvez, 2007).

### Inorganic carbon delivery components are conserved between *Chlorella* and *Chlamydomonas*, but pyrenoid structural components are not

We next set out to identify the protein components involved in Ci delivery and pyrenoid assembly in *Chlorella* pCCM. In *Chlamydomonas*, Ci uptake at HC is predominantly via passive CO_2_ diffusion to Rubisco dispersed throughout the chloroplast stroma (60% outside the pyrenoid) (Wang, Stessman and Spalding, 2015; Mackinder et al., 2016). Under VLC conditions, large transcriptomic (Fang et al., 2012; Brueggeman et al., 2012) and proteomic changes (Strenkert et al., 2019) are linked to active Ci uptake in *Chlamydomonas*. The high affinity for bicarbonate uptake and the formation of a pyrenoid under LC and VLC in *Chlorella* suggest that CCM-associated proteins may also be enriched under these conditions. To identify candidate CCM components, we performed a comparative proteomic analysis of cells grown in VLC and HC (Supplementary table 1). Using a log2 fold-change threshold of >2 (>4-fold enrichment) and a -log_10_ adjusted p-value threshold of >1.3 (p_adj < 0.05), we identified a total of 256 proteins above these cut-offs at VLC vs. HC (Supplementary Table 2). Among these significantly enriched proteins, 78 of them showed sequence similarity (BLAST) with proteins in *Chlamydomonas* proteome (Supplementary Table 3 and 4). As expected, top hits from this analysis included known CCM components such as HLA3, CAH2, CAH3, LCIB/LCIC, BSTs, and the CCM regulator CAS (Fig. 1e; Supplementary Table 2). We also observed upregulated homologs of mitochondrial proteins that are linked to CCM function in *Chlamydomonas*, including CAH4/5 and CCP1/2 (Fig. 1e; Supplementary Table 1 and 2). Together, these results indicate the conservation of Ci delivery machinery in *Chlorella* and *Chlamydomonas*.

Outside of Ci transport and regulatory components, we saw a significant increase in CsLinker (CSI2_123000012064) under VLC conditions, the functional analogue to EPYC1 that has minimal sequence conservation. However, both in our mass spectrometry data and through protein sequence BLAST, we saw no homologs of known *Chlamydomonas* pyrenoid structural components such as SAGA1, SAGA2, MITH1, and RBMP2. Our data indicates that Ci delivery machinery is conserved across distant Chlorophyte species (Fig. 1f) but that pyrenoid structural components are not. This supports the hypothesis that pyrenoids convergently evolved within the green lineage (Long, Matsuda and Moroney, 2024; Barrett et al., 2024, 2026) after the split of the core Chlorophyte classes.

### Cryo-electron tomography reveals the native architecture of the *Chlorella* pyrenoid

The apparent lack of conservation of pyrenoid structural components between *Chlorella* and *Chlamydomonas* inspired us to gain a better understanding of the architecture of the *Chlorella* pyrenoid. Since TEM data from our work and previous studies (Carfagna et al., 2013; Bai et al., 2019; Guo et al., 2022) revealed distinct pyrenoid morphology in *Chlorella* compared to *Chlamydomonas*, we turned to cryo-ET to resolve native molecular details. Cryo-ET has proven extremely powerful for visualizing pyrenoid architecture (Engel et al., 2015; Shimakawa et al., 2024; Nam et al., 2024), analyzing pyrenoid protein organization (Kumar et al., 2026; Freeman Rosenzweig et al., 2017), and discovering pyrenoid protein components (Shimakawa et al., 2024). We performed *in situ* cryo-ET on LC-adapted cells that have a fully formed pyrenoid matrix encapsulated by a starch-sheath.

In tomograms of LC-adapted cells, the pyrenoid displayed the typical architecture of an elliptical Rubisco matrix, flanked on both sides by starch plates and bisected by a central pair of appressed thylakoids traversing the long axis of the matrix (Fig. 2a,b). Along these PTTs, we observed two notable, previously unreported architectural features. First, in several regions the thylakoid membranes locally constrict, forming channel-like fenestrations that connect the Rubisco matrix with the stromal space between the two PTTs (Fig. 2a; Extended Data Fig. 4). These holes thereby create a continuous path for diffusion between the Rubisco matrix and the surrounding chloroplast stroma outside the pyrenoid, perhaps serving a similar function to the minitubules seen in *Chlamydomonas* (Engel et al., 2015). Second, segments of the PTT pair adopted a twisted, braid-like arrangement at multiple positions along their length (Fig. 2c, panel iv). To our knowledge, this membrane morphology has not been described previously for any PTTs.

**Figure 2:**
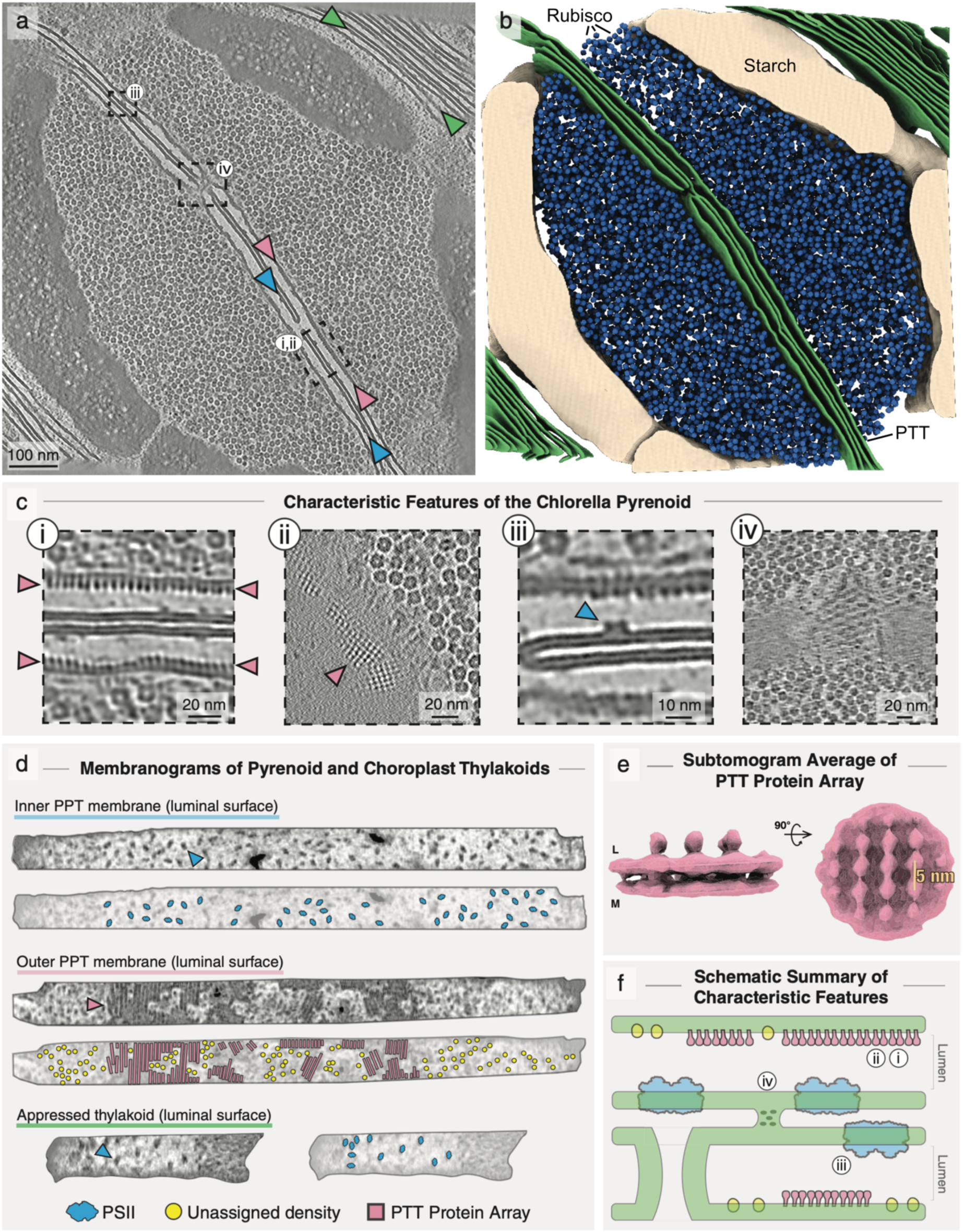
Cryo-ET reveals the ultrastructural organization and distinctive molecular features of the *Chlorella* pyrenoid. a) Central tomographic slice through the pyrenoid showing the Rubisco condensate, two appressed pyrenoid traversing thylakoids (PTT), and surrounding starch plates. Dashed boxes correspond to features in (c), membranes between arrowheads correspond to membranograms indicated with the same color in (d). b) Segmentation of tomogram in (a), showing major pyrenoid components: thylakoid membranes (green), starch plates (beige), and Rubisco matrix (blue). c) Enlarged views from (a) highlighting: (i) side view of the PTT protein array (pink arrowhead), (ii) orthogonal membrane view of the PTT protein array revealing its square lattice organization, (iii) close-up of a PSII complex (blue) within a PTT membrane, and (iv) a braided-like membrane morphology suggestive of a distinct membrane phase or protein structure in the PTT. d) Membranograms depicting tomogram densities projected on membrane luminal surfaces, corresponding to membranes between arrowheads of the same color in (a). A second view of each membranogram is shown with annotated particles. PSII complexes (blue) are found in appressed thylakoids (both internal-facing PTT membranes and thylakoids outside the pyrenoid); Rubisco-facing PTT membranes contain protein arrays (pink) and small densities that were not assigned an identity (yellow). e) Subtomogram average of PTT arrays, exhibiting clear membrane connections and a 5 nm lattice repeat of globular densities extending into the lumen. Isosurface renderings shown from side view (membrane cross-section; L: thylakoid lumen and M: Rubisco matrix) and top view (orthogonal to membrane). See also Extended Data Fig. 5. f) Schematic summarizing characteristic structural features of the *Chorella* PTT pair.

To examine the protein complexes embedded in the PTT membranes, we generated membranograms to visualize densities extending from each membrane’s luminal surface (Yamauchi et al., 2024; Wietrzynski et al., 2020; Yan et al., 2025) (Fig. 2a,c,d). This analysis revealed differential protein occupancies between the outer bilayers that face the Rubisco matrix and the inner bilayers that are closely appressed to the other thylakoid of the PTT pair. On the inner membranes, densities consistent with Photosystem II (PSII) complexes were observed projecting into the thylakoid lumen (Fig. 2a,c,d; blue), similar in appearance and distribution to PSII in appressed thylakoids outside the pyrenoid (Fig. 2a,d; green). In contrast, the Rubisco-facing membrane contained local patches of tightly packed protein densities arranged in arrays that span the bilayer and extend into the PTT lumen (Fig. 2a,c,d; pink). In addition, punctate luminal densities of unknown identity were observed surrounding these arrays (Fig. 2d; yellow).

We hypothesize that the array densities may correspond to carbonic anhydrases, as luminal PTT-localized carbonic anhydrases are key components of both diatom and *Chlamydomonas* CCMs (Sinetova et al., 2012; Karlsson et al., 1998; Shimakawa et al., 2023). To further test this hypothesis, we performed subtomogram averaging of the array-forming densities (Fig. 2e; Extended Data Fig. 5). The resulting average is consistent with 5 nm diameter globular repeats projecting into the lumen, spaced 5 nm apart to form a square lattice on the membrane. Although these densities cannot be unambiguously assigned at the obtained resolution, AlphaFold-predicted CsCAH3 fits within the map dimensions, supporting CsCAH3 as a candidate component of the luminal array (Extended Data Fig. 5) and consistent with its increased abundance under limiting CO_2_ conditions (Fig. 1e; Supplementary Table 2). Together, these observations highlight the distinct architecture of PTT sheets in *Chlorella* (Fig. 2f).

### Multiple pyrenoid proteins contain a shared Rubisco binding motif

Although cryo-ET analysis of pyrenoid ultrastructure revealed multiple distinct features compared to *Chlamydomonas*, some aspects are similar, including the presence of starch-matrix and membrane-matrix interfaces. As discussed, pyrenoid proteins essential to these interfaces in *Chlamydomonas* (e.g., MITH1, RBMP2, SAGA1, SAGA2) do not have homologs in *Chlorella*. In *Chlamydomonas*, these proteins contain structural or functional domains fused to Rubisco-binding motifs (RBMs) that have high sequence homology with the matrix linker protein, EPYC1 (Meyer et al., 2020). We therefore hypothesized that we might find analogous proteins in *Chlorella* harboring RBMs similar to those found in the independently evolved linker protein of *Chlorella*, CsLinker.

To search for candidates, we performed BLAST of the CsLinker RBM against the *Chlorella* proteome. In addition to CsLinker, three proteins were found to contain CsLinker-like Rubisco-binding regions (CSI2_123000008029, CSI2_123000003237 and CSI2_123000010418). CSI2_123000008029 is a homolog of CsLinker that contains no additional structural domains, but is expressed to a very low level under LC conditions (<1% of CsLinker). The remaining two proteins contained multiple RBMs and intrinsically disordered regions (IDRs), similar to CsLinker but with additional functional or structural domains (Fig. 3a, Extended Data Fig. 6).

**Figure 3.**
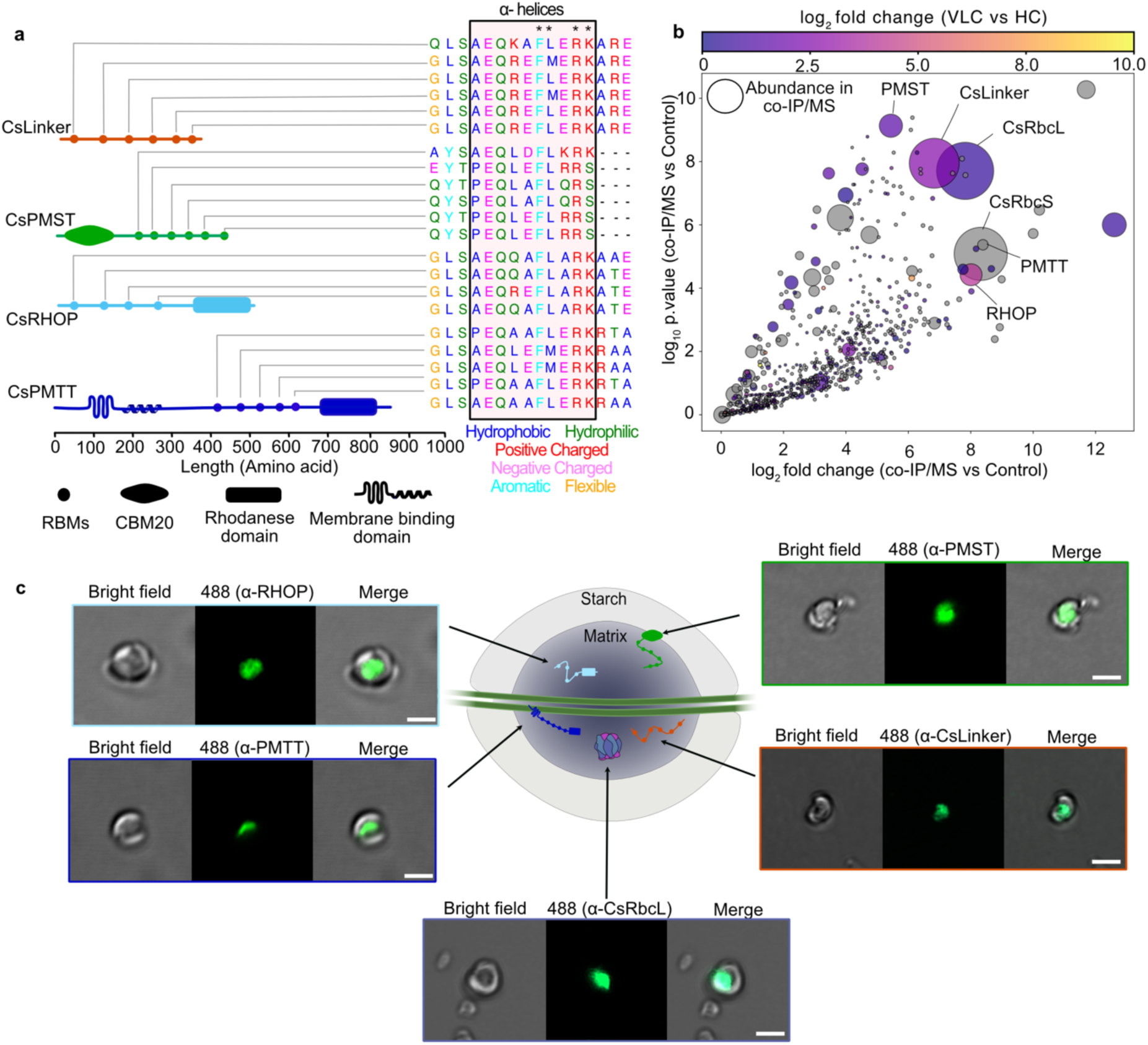
Identification of *Chlorella* pyrenoid proteins based on RBMs and validation of their interaction with CsRubisco. a) The known pyrenoid protein CsLinker contains six RBMs. Similar RBMs were identified in three additional *Chlorella* proteins: CsPMST, CsRHOP, and CsPMTT. The key residues for RbcL interaction are shown with *. b) co-IP/MS using CsRubisco as bait confirms enrichment of all three candidate proteins. The colorbar shows proteins that have log_2_ fold change of VLC vs. HC > 0. All the proteins that are not significantly enriched at VLC are shown in gray. The size of data points in the plot depicts the relative abundance of the protein in the co-IP/MS. co-IP/MS data replotted from Barrett et al. (2024). c) Immunofluorescence of isolated *Chlorella* pyrenoids reveals distinct localization: CsPMTT at traversing thylakoids, CsPMST at starch plates, and CsRHOP together with CsLinker within the pyrenoid matrix. Scale bar, 2 μm.

The proteins were named based on their putative roles in pyrenoid function, inferred from the presence of additional domains. CSI2_123000003237 has a transmembrane domain (TMD) at the N-terminal, a largely disordered region with 5 RBMs in the middle and a Rhodanese domain at the C-terminal. We propose it may mediate interactions between Rubisco and PPTs, and we therefore name it Putative Matrix-Thylakoid Tether (PMTT) (Extended Data Fig. 7). A Rhodanese domain was predicted at the C-terminus of PMTT; however, this domain lacked the catalytic cysteine at its active site, suggesting that this domain may be inactive (Extended Data Fig. 8). CSI2_123000010418 harbors a predicted active Rhodanese domain (Extended Data Fig. 8) in addition to RBMs, but unlike PMTT lacks a TMD suggesting it may be soluble. We name it RHOdanese containing Pyrenoid protein (RHOP). In these searches we did not identify a homolog of SAGA1 or SAGA2, which would contain a CsLinker-like RBM as well as a carbohydrate-binding module 20 (CBM20) domain for starch interaction (Ngo et al., 2019). To search for homologous proteins, we instead used a BLAST search of the CBM20 domain of SAGA1. Using this approach, we identified nine proteins, six of which contained an N-terminal CBM20 in addition to a largely disordered C-terminus. By aligning the Rubisco-binding region of CsLinker against these hits, we identified a single protein, CSI2_123000007310, with sequence homologous to the RBM of CsLinker. This protein, which we name Putative Matrix-Starch Tether (PMST), contains six RBMs, which differ slightly from the other motifs found in CsLinker, PMTT and RHOP, but retain key Rubisco-binding residues (Fig. 3a) (Barrett et al., 2024).

### PMST, RHOP, and PMTT are Rubisco-interacting pyrenoid proteins with different sub-pyrenoid localizations

We initially sought to validate the interaction of PMST, RHOP, and PMTT with Rubisco. We re-analyzed the CsRubisco co-immunoprecipitation coupled with mass spectrometry (co-IP/MS) data that we previously used to identify CsLinker (Barrett et al., 2024). Along with RbcL, RbcS and CsLinker, PMST, RHOP, and PMTT were all significantly enriched in the CsRubisco co-IP/MS pulldown (Fig. 3b). PMST and RHOP were also significantly more abundant at VLC compared to HC, whereas PMTT was only marginally enriched (Fig. 3b and Supplementary Tables 1 and 2).

To study the localization of these proteins to the pyrenoid and investigate their roles in pyrenoid function, we attempted to develop a reliable transformation protocol for *Chlorella*, but were unsuccessful. Consequently, we conducted immunofluorescence analysis of isolated pyrenoids using specific antibodies raised against specific peptides of each protein (Barrett et al., 2024). CsLinker and RHOP had a localization pattern that was similar to RbcL, consistent with them being components of the pyrenoid matrix. Although PMST contains a CBM20 domain, immunofluorescence analysis showed it localizing throughout isolated pyrenoids. This may reflect the collapsed nature of isolated pyrenoids which produces a fluorescence signal that appears to emanate from throughout the pyrenoid rather than at a discrete starch surface. PMTT had a much narrower signal distribution which was positioned consistent with the location of the thylakoids that traverse the matrix (Fig. 3c).

### PMST, RHOP, and PMTT condense CsRubisco in vitro but not efficiently at physiological concentrations

Since PMST, RHOP, and PMTT contain predicted disordered regions with multiple RBMs similar to CsLinker, we tested whether they could also facilitate the liquid-liquid phase separation (LLPS) of CsRubisco (Barrett et al., 2024). We mixed recombinant PMST, RHOP and PMTT purified from *Escherichia coli* with CsRubisco purified from *Chlorella* (Extended Data Fig. 9). All three proteins could drive LLPS of CsRubisco, forming spherical droplets comparable to droplets formed by mixing CsLinker with CsRubisco (Fig. 4a) (Barrett et al., 2024). This raised the question of whether these proteins are at physiologically relevant concentrations to drive the LLPS of CsRubisco in vivo. To address this, we carried out absolute quantification mass spectrometry using standard curves of recombinantly produced PMST, RHOP, and PMTT (Fig. 4b; Extended Data Fig. 10; Supplementary Table 5) (Barrett et al., 2024). Based on the reported chloroplast volume for *Chlorella pyrenoidosa* (Guo et al., 2022), we estimated the concentration of PMST, RHOP, and PMTT to be approximately 0.657 ± 0.12 μM s.d., 0.626 ± 0.08 μM s.d., and 2.42 ± 0.52 μM s.d., respectively. The CsLinker and CsRubisco concentrations were previously estimated to be 6.24 ± 0.73 μM s.d. and 2.95 ± 0.06 μM s.d., respectively (Barrett et al., 2024).

**Figure 4.**
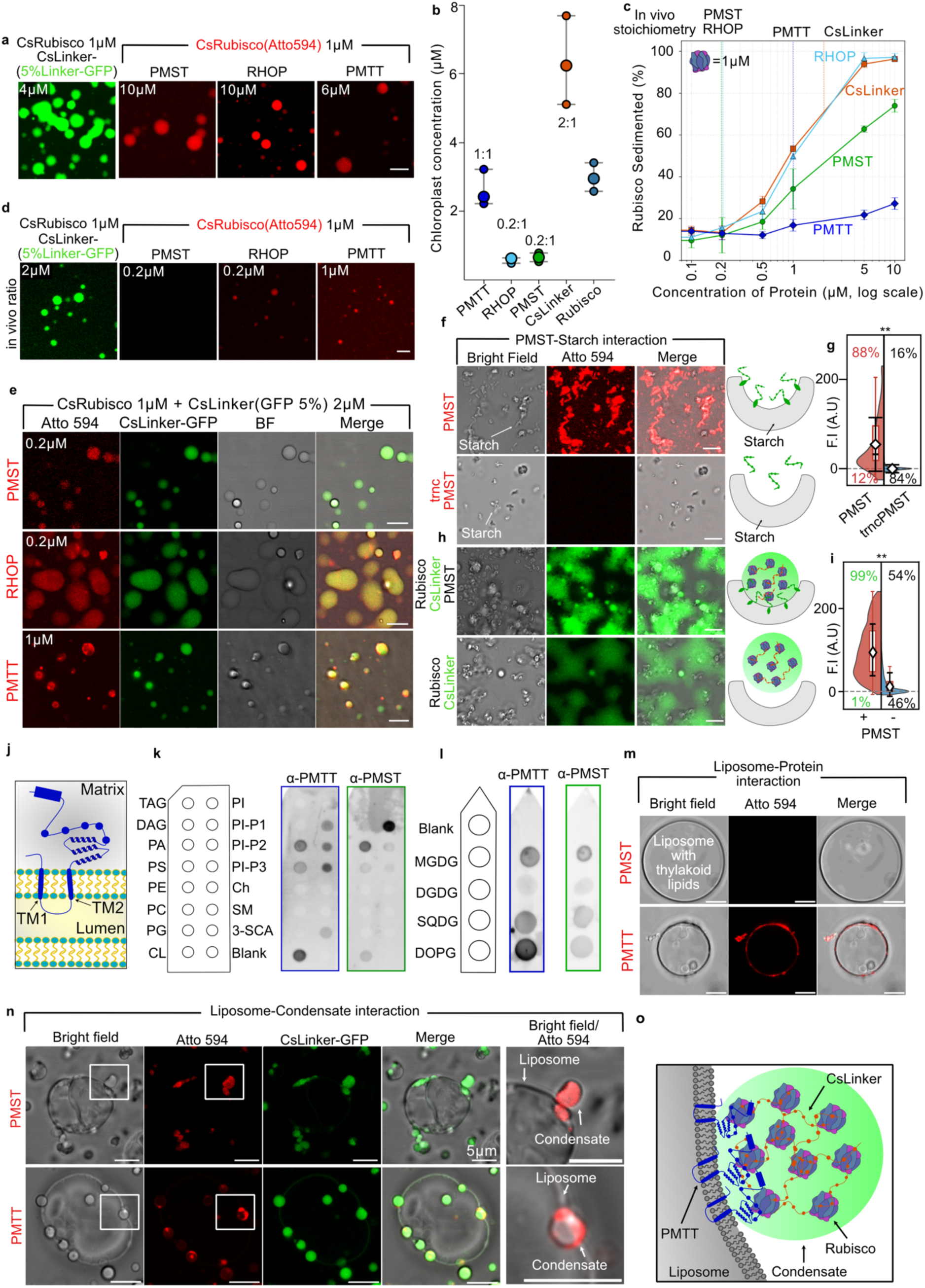
Biochemical characterization of identified pyrenoid proteins based on predicted functional domains. a) In vitro condensate formation by CsLinker, PMST, RHOP and PMTT when mixed with CsRubisco at the stated concentrations. Scale bar, 5 μm. b) Absolute in vivo quantification of *Chlorella* pyrenoid proteins (n=3). Ratios are relative to Rubisco. c) Rubisco sedimentation assays performed with *Chlorella* pyrenoid proteins at increasing concentrations. Rubisco concentration was fixed at 1 μM, as determined from two replicate experiments. The corresponding in vivo protein concentrations, derived from absolute quantification, are indicated at the top. d) At ratios similar to in vivo, PMST, RHOP and PMTT exhibited weak phase separation. Scale bar, 5 μm. e) Multi-component assays with CsRubisco, CsLinker, and individual candidate proteins at in vivo stoichiometries promotes efficient condensate formation, with candidate proteins colocalizing to condensates. Scale bar, 5 μm. f) Atto594-labelled PMST localizes specifically to starch plates isolated from *Chlorella*, whereas localization is lost upon CBM20 removal. Scale bar, 5 μm g) Raincloud plot quantifying the colocalization of fluorescence signal with starch granules from the image (f). In the analysed frame, 88% of starch molecules exhibited colocalization with fluorescence signal from full-length PMST above the set threshold, compared with only 16% for the CBM20-truncated PMST construct. F.I is the fluorescence intensity and A.U is arbitrary units. Statistical comparison: Welch’s unpaired two-tailed t-test; **p < 0.05. h) Condensates formed from CsRubisco, CsLinker, and PMST associate with starch plates, whereas those lacking PMST do not. Scale bar, 5 μm i) Raincloud plot quantifying colocalization between GFP fluorescence (from CsLinker) and starch granules from the images shown in (h). Starch–fluorescence colocalization was 99% when PMST was present within the condensate, compared with 54% when PMST was absent. F.I is the fluorescence intensity and A.U is arbitrary units. Same statistics as in (g). j) Schematic of PMTT showing the predicted transmembrane (TM) domain anchoring to membranes and extending into the pyrenoid matrix. k) LPO assays with PMTT and PMST using commercially available lipid strips. l) LPO assays using PMTT and PMST with in-house prepared lipid strips containing thylakoid membrane lipids. m) Fluorescence microscopy of Atto594-labelled PMTT with giant unilamellar vesicle (GUV) liposomes composed of thylakoid-like lipids. PMST was used as a negative control. Scale bar, 5 μm n) Multi-component condensates containing Rubisco, CsLinker, and Atto594-labelled PMTT or PMST in presence of GUVs show PMTT localization to the membrane-condensate interface. Scale bar, 5 μm o) Schematic of PMTT localizing to the membrane-condensate interface in an in vitro setting.

To determine the concentration required to efficiently demix a physiologically relevant concentration of CsRubisco, we performed titration sedimentation assays at a fixed CsRubisco concentration of 1 μM (Extended Data Fig. 11). RHOP and CsLinker had comparable efficiency, requiring greater than 1 µM but less than 5 µM to reach maximum partitioning of CsRubisco into the dense phase (Fig. 4c). PMST was less efficient, with maximum partitioning not achieved at 10 µM of PMST (Fig. 4c). Finally, PMTT had much lower efficiency, with <30% of Rubisco sedimented at 10 µM PMTT (Fig. 4c). These results support that, apart from CsLinker (Barrett et al., 2024), the in vivo concentrations of the candidate proteins are insufficient to reach maximum CsRubisco partitioning into the dense phase. Consistent with this, microscopy-based condensation assays revealed that at in vivo-like stoichiometries, only CsLinker efficiently promoted condensation of Rubisco into large droplets (Fig. 4d).

### PMST, RHOP, and PMTT are recruited to CsLinker-CsRubisco condensates

We next tested whether PMST, RHOP, and PMTT could be recruited to condensates that had been formed with CsLinker and CsRubisco at physiologically relevant concentrations. For this, we performed condensation assays in which CsRubisco, CsLinker, and either PMST, RHOP or PMTT were mixed at in vivo ratios. To visualize recruitment, 5% of CsLinker was GFP-tagged, while each candidate protein was fluorescently labelled with the red fluorescent dye Atto594. Under these conditions, all three candidate proteins localized to the CsRubisco-CsLinker condensates (Fig. 4e). Interestingly, while PMST and RHOP partitioned homogeneously throughout the condensate, PMTT exhibited a heterogeneous distribution, characterized by enrichment at the periphery of the condensate or localized clustering within the condensate (Fig. 4e). We hypothesize that this behavior is driven by the N-terminal transmembrane domain (TMD) as this represents the main difference between PMTT and RHOP, the latter of which localized uniformly within the condensate.

These findings support that RBMs fused to structural or enzymatic domains enable the recruitment of proteins to Rubisco condensates, similar to that observed for RBM-containing pyrenoid proteins in *Chlamydomonas* (Meyer et al., 2020). This recruitment could enable partitioning of biochemical processes, regulate condensate properties, enable pyrenoid assembly and/or define pyrenoid architecture. As PMST and PMTT are fused with domains that could interact with starch and membranes, respectively, we next further explored their functional roles.

### PMST mediates condensate-starch interaction

PMST harbours a predicted N-terminal CBM20 starch binding domain fused to an IDR containing six RBMs (Fig. 3a). To validate the starch interaction of this protein, we recombinantly produced full-length PMST and a truncated version of PMST lacking the CBM20 domain (trncPMST). Both PMST and trncPMST were labelled with Atto594 and incubated with starch granules extracted from *Chlorella*. Quantification of colocalization between starch granules and the Atto594 signal via confocal microscopy revealed that 88% of starch granules exhibited Atto594 localization with PMST, whereas this proportion was reduced to 16% in the case of trncPMST (Fig. 4f,g; Extended Data Fig. 12). Whilst CBM20 domains are primarily associated with starch binding, previous work has suggested that this domain may directly disrupt the surface of starch (Southall et al., 1999). To check if PMST affects starch granule morphology, we incubated isolated starch from *Chlorella* with PMST and imaged using scanning electron microscopy (SEM); however, no notable structural differences were observed compared to untreated control starch suggesting that PMST alone does not impact starch morphology (Extended Data Fig. 13).

Given that PMST interacts with Rubisco, is recruited to Rubisco-CsLinker condensates, and binds to starch, we hypothesised that its functional role in vivo could be to mediate the interface between the Rubisco matrix and the surrounding starch plates. We tested this in vitro by assembling multicomponent condensates composed of Rubisco, CsLinker (5% CsLinker-GFP), and PMST at physiological concentrations and incubating them with starch granules. Colocalization analysis of the GFP signal from the condensate and starch granules showed that 99% of starch granules exhibited GFP signal in the presence of PMST (Fig. 4h,i). In contrast, only 54% of starch granules showed GFP localization in its absence, confirming enhanced starch–condensate interaction in the presence of PMST (Fig. 4h,i; Extended Data Fig. 14). These observations indicate that the CBM20 domain mediates PMST localization to starch plates. Furthermore, the targeting of PMST-containing condensates to starch suggests that PMST may contribute to the assembly or organization of starch around the pyrenoid matrix in vivo.

### PMTT interacts with membrane lipids in vitro

The presence of a transmembrane domain in PMTT (Fig. 4j) and its localization to the PTTs (Fig. 3c) suggest a potential role in recruiting thylakoid membranes into the pyrenoid matrix or stabilizing the pyrenoid matrix around the PTTs.

To validate the membrane interaction capability of PMTT, we performed a Protein-Lipid Overlay (PLO) assay using a commercial lipid strip containing a diverse array of lipids with varying physicochemical properties (Fig. 4k). Given that PMTT is a basic protein with an isoelectric point (pI) of 9.40, we utilized PMST, that has a comparable pI of 9.10, as a control to distinguish specific binding events from those arising from electrostatic attractions. Neither PMTT or PMST showed detectable interaction with neutral lipids, including triacylglycerol (TAG), diacylglycerol (DAG), phosphatidylethanolamine (PE), phosphatidylcholine (PC), cholesterol (Ch), and sphingomyelin (SM) (Fig. 4k). This absence of binding excludes non-specific hydrophobic partitioning and suggests that an electrostatic trigger could be a prerequisite for membrane recruitment. Both PMTT and PMST showed varying levels of interaction with negatively charged lipids of Phosphatidylinositol (PtdIns) family (PI-P1, P2 and P3), Phosphatidylserine (PS) and Phosphatidic acid (PA), which may arise from the electrostatic interaction of the proteins with these lipids. PMTT showed strong interaction with Cardiolipin (CL) found in mitochondrial membranes and 3-Sulfogalactosylceramide (3-SCA) carrying a sulfated sugar headgroup similar to sulfoquinovosyldiacylglycerol (SQDG) found in thylakoid membranes.

Next, to explore the binding of PMTT to thylakoid relevant lipids, we generated a custom PLO array containing the four major thylakoid lipids: monogalactosyldiacylglycerol (MGDG), digalactosyldiacylglycerol (DGDG), SQDG, and dioleoyl-PG (DOPG) (Fig. 4l) (Douce and Joyard, 1990). On this native array, PMTT showed clear enrichment on MGDG, SQDG, and DOPG compared to the PMST control. Although both MGDG and DGDG are neutral galactolipids, PMTT preferentially bound MGDG (Fig. 4l). MGDG is a non-bilayer, cone-shaped lipid known to induce membrane curvature, whereas DGDG is a bilayer-forming cylindrical lipid; this preference suggests PMTT may specifically localize to curved membrane regions or enrich the curvature inducing lipids at bound sites (Simidjiev et al., 2000; Hoyo, Guaus and Torrent-Burgués, 2016; Koldsø and Sansom, 2012). The robust binding to SQDG confirms the affinity for sulfated lipids suggested by the initial 3-SCA result. Whilst PLOs provide a starting point for screening protein-lipid interactions they do not fully mimic the fluid, two-dimensional bilayer of a membrane, we therefore turned to liposome-based assays.

### PMTT interacts with liposomes and potentially tethers them to condensates

To validate the interaction of PMTT with thylakoid-like membranes, we generated liposomes mimicking thylakoid lipid composition (MGDG:DGDG:SQDG:DOPG=5:3:1:1) (Extended Data Fig. 15) (Mizusawa and Wada, 2012). We employed fluorescence microscopy using Atto594-labelled PMTT, which demonstrated clear localization of the protein to the liposomes (Fig. 4m). Atto594-labelled PMST showed no liposome interaction, confirming the specificity of PMTT towards native membranes (Fig. 4m).

Finally, we assessed the ability of PMTT to tether these thylakoid-like liposomes to Rubisco-CsLinker condensates, with PMST used as a control. We saw clear association of condensates with membranes for both PMTT and PMST. Whilst the quantification of condensate attachment to liposomes is challenging due to variations in liposome and condensate number and size between experiments, we made two key observations: 1) Condensate attachment to liposomes in the presence of PMTT typically resulted in the flattening or negative curvature of the membrane (Fig. 4n,o; Extended Data Fig. 16); 2) We saw an enrichment of PMTT at the condensate-liposome interface, whereas PMST maintained a uniform distribution throughout the condensate (Fig. 4n, Extended Data Fig. 16). These observations support that PMTT is enriched to the condensate-liposome interface and impacts membrane curvature. Condensation on the liposome surface could provide a compressive force leading to curvature (Mondal and Baumgart, 2023), or enriched PMTT at the liposome-condensate boundary could be recruiting specific lipids that favour negative curvature. Taken together, PMTT interacts with negatively charged and thylakoid-specific lipids. The ability of PMTT to associate with liposomes containing native lipid compositions, combined with our previous immunofluorescence data (Fig. 3c), confirms its identity as a membrane-bound protein localized to the pyrenoid. Based on the condensate-liposome experiments, where PMTT localized to the interface and potentially induced inward curvatures, we propose that PMTT may function as an anchoring protein that tethers PTT membranes to the Rubisco matrix.

### *Chlorella* pyrenoid proteins exhibit domain-specific localization in *Chlamydomonas*

To investigate the in vivo functionality of *Chlorella* pyrenoid proteins, we employed heterologous chloroplast expression in *Chlamydomonas*. The RBMs in CsLinker bind to a sequence conserved surface of the Rubisco large subunit (RbcL). Owing to this sequence conservation, CsLinker also interacts with the *Chlamydomonas* Rubisco large subunit (CrRbcL) and has been shown to drive the condensation of *Chlamydomonas* Rubisco (CrRubisco) in vitro and in vivo (Barrett et al., 2024). Because the key Rubisco interacting residues are conserved across the RBMs in CsLinker and the three candidate proteins, we hypothesized that these proteins would similarly interact with *Chlamydomonas* CrRubisco through binding to its large subunit.

In line with our immunofluorescence localization data, RHOP, fused to an N-terminal mCherry, localized to the pyrenoid matrix (Fig. 5a, Extended Data Fig. 17) and PMTT fused to a C-terminal mVenus localized to the center of the pyrenoid, where the thylakoid-derived pyrenoid tubules converge to form a reticulated “knot-like” structure (Fig. 5b, Extended Data Fig. 17). In *Chlamydomonas*, several proteins critical for pyrenoid structure and function, such as SAGA1 and MITH1, have also been reported to localize to the pyrenoid tubules (Meyer et al., 2020; Hennacy et al., 2024). MITH1 and SAGA1 were recently identified as facilitators of tubule extension from the thylakoids into the Rubisco matrix, with *mith1* knockout strains lacking traversing tubules. Based on the similar sub-pyrenoid localization patterns, we hypothesized that PMTT might functionally complement MITH1. However, attempts to complement the *mith1* mutant using the PMTT-mVenus construct failed, with no detectable mVenus signal (Extended Data Fig. 18b). Although not further explored, the lack of detectable expression via confocal microscopy could be due to the absence of pyrenoid tubules in the *mith1* mutant, resulting in either a diffuse PMTT signal throughout the thylakoid network or degradation of the protein.

**Figure 5:**
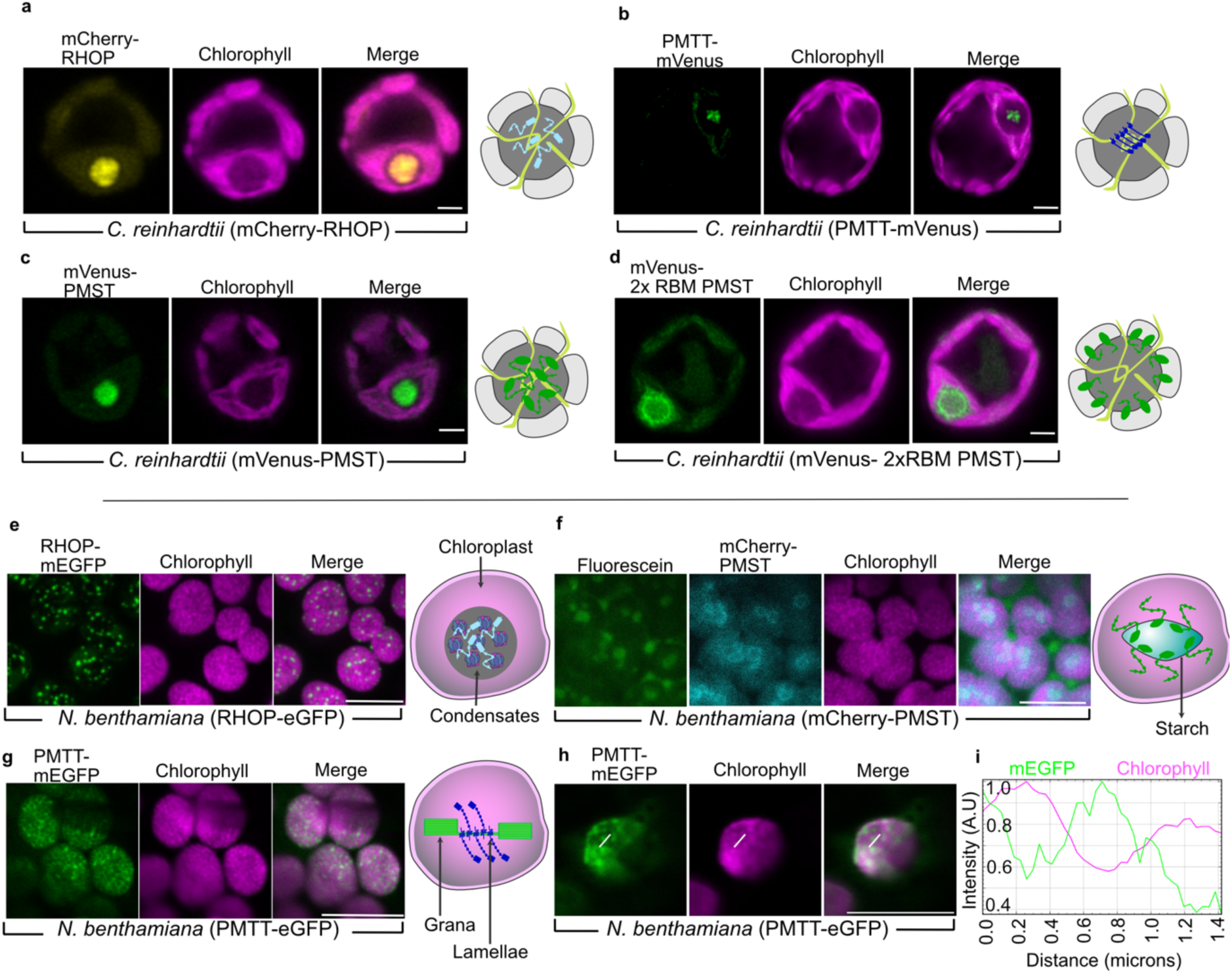
In vivo localization of *Chlorella* pyrenoid proteins in *Chlamydomonas*. (a–e) and in the vascular plant *Nicotiana benthamiana* (f–i). a) CsRHOP localizes to the pyrenoid matrix. b) CsPMTT localizes to the center of the pyrenoid matrix, coinciding with the “central knot” of reticulated membrane tubules. c) Full-length CsPMST localizes to the pyrenoid matrix, whereas d) truncated CsPMST containing only two RBMs localizes to the matrix–starch interface. e) Expression of CsRHOP in *N. benthamiana* chloroplasts yields small puncta. f) Chloroplast targeting of CsPMST achieved by replacing the native signal peptide with a plant chloroplast signal peptide; fluorescein labeling confirming localization to starch granules (Extended Data Fig. 19). g) CsPMTT localization in mesophyll chloroplasts shows mEGFP signals enriched in chlorophyll-depleted gaps i) CsPMTT localization in epidermal chloroplast. Plot of normalized fluorescence intensity values through a cross section representing two grana stacks. Scale bar, 10 μm.

Based on the presence of a starch-binding domain (CBM20), we hypothesized that PMST would localize to the matrix–starch interface, as observed for *Chlamydomonas* proteins SAGA1 and SAGA2 (Meyer et al., 2020). However, PMST with a C-terminal mVenus tag localized predominantly to the pyrenoid matrix (Fig. 5c). We speculated that this unexpected localization was due to strong interactions between the six RBMs on PMST and the large subunit of Rubisco in the absence of competition with CsLinker for the binding sites that would be present in a wild-type *Chlorella* pyrenoid. To test this, we generated PMST variants containing one to four RBMs. PMST proteins with at least four RBMs consistently localized to the pyrenoid matrix, while those with one to three RBMs localized to the matrix–starch interface (Fig. 5d, Extended Data Fig. 17). This observation suggests that RBM number most likely is finely tuned to ensure correct localization and potentially functionality.

Considering that PMST contains a CBM20 domain and, like SAGA1 in *Chlamydomonas*, localizes to the periphery of the pyrenoid matrix (as observed for truncated PMST), we tested whether PMST or its truncated variants could complement the *saga1* mutant that has multiple pyrenoids. Whilst PMST protein variants expressed in *saga1*, none of them resulted in single pyrenoid formation nor rescued the *saga1* mutant phenotype in VLC (Extended Data Fig. 18).

In summary, although PMTT and PMST localize to their predicted sub-pyrenoid domains when exogenously expressed in *Chlamydomonas*, PMTT fails to complement the *mith1* mutant that lacks pyrenoid tubules, and PMST fails to complement *saga1* that has a starch-associated pyrenoid defect. This suggests that while these proteins may serve analogous roles in the two evolutionarily distant species of green algae, they are not cross-compatible.

### *Chlorella* pyrenoid proteins localize to distinct regions of the *Nicotiana benthamiana* chloroplast

Engineering a pyrenoid-based CCM into C3 crops promises to be a transformative strategy for enhancing photosynthetic efficiency (Fei et al., 2022; Long, Marshall-Colon and Zhu, 2015; Long et al., 2025). The binding of *Chlorella* RBMs to plant Rubisco makes engineering a *Chlorella*-based pyrenoid a feasible prospect that could be universally applied across plant species (Barrett et al., 2024). Our identification and characterization of the *Chlorella* pyrenoid proteins significantly expands the toolbox for this endeavour. As a primary step toward validating these new components for plant engineering, we transiently expressed them in *Nicotiana benthamiana* mesophyll cells.

RHOP with a C-terminal mEGFP tag localized to the chloroplast and formed discrete puncta, indicating potential condensation of Rubisco in planta (Fig. 5e). This is consistent with the in vitro LLPS propensity of RHOP, which closely resembles that of CsLinker, also previously shown to form condensates in *N. benthamiana* chloroplasts (Barrett et al., 2024). PMST did not localize to the chloroplast with its native signal peptide. Replacing the native peptide with the plant chloroplast signal peptide derived from AtRbcS1A successfully directed PMST into the chloroplast (Extended Data Fig. 19). To examine its association with starch granules, PMST was tagged with mCherry and starch was stained with fluorescein, revealing PMST enrichment at the periphery of starch within the chloroplast (Fig. 5f). PMTT with a C-terminal mEGFP tag localized to the chloroplast, with the mEGFP signal consistent with association to *N. benthamiana* thylakoids. This localization is similar to algal proteins previously shown to be localized to Arabidopsis stromal lamellae, where the peak GFP signal is interspersed with chlorophyll fluorescence (Fig. 5g,h,i) (Adler et al., 2024).

### Perspective

Pyrenoids are globally important biomolecular condensates with large engineering potential to enhance photosynthesis (Long et al., 2025). Their convergent evolution means that their molecular composition and emergent architecture cannot be inferred across clades. To advance our understanding of pyrenoid evolution, structure, function and engineering potential, we have integrated diverse methodologies to understand carbon fixation in *Chlorella*, a representative of the Trebouxiophyceae algae – a clade in which very little is known regarding CCM and pyrenoid function. We show that *Chlorella* has an inducible CCM that is coordinated with large proteomic and structural changes, with similarities to those observed in *Chlamydomonas*. These include upregulation of inorganic carbon uptake machinery that is largely conserved with that of *Chlamydomonas*, the recruitment of Rubisco to the pyrenoid, and the formation of a Rubisco matrix-encapsulating starch-sheath at limiting CO_2_. However, there are key differences with *Chlamydomonas* in pyrenoid architecture and the underpinning pyrenoid structural components, providing compelling evidence for the convergent evolution of pyrenoid organization and its constituent proteins, even within relatively closely related green algal clades.

### Architectural and structural insights into the *Chlorella* pyrenoid

In-depth analysis of native pyrenoid architecture using cryo-ET revealed notable differences from *Chlamydomonas*, including two appressed thylakoid sheets (PTTs) that bisect the Rubisco matrix and surrounding starch sheath. PTTs with similar PTT morphology have been reported in the pyrenoids of diatoms (Nam et al., 2024; Shimakawa et al., 2024). In contrast, the multiple pyrenoid traversing membranes in *Chlamydomonas* undergo structural remodelling as they enter the pyrenoid, forming cylindrical tubules that enclose minitubules. These tubules then extend towards the centre of the pyrenoid where they converge to form a reticulated central knot (Engel et al., 2015). In the *Chlorella* PTTs we could confidently assign PSII to the two appressed inner membranes of the PTT pair, which has not been reported in the pyrenoid tubules of *Chlamydomonas*. Densities were not observed that could be attributed to other photosynthetic electron transport chain components, although we cannot strictly exclude this possibility. As is the case in conventional thylakoids, PSII and its associated LHCII antennae most likely mediate the stacking between the pair of PTTs (Daum et al., 2010; Wietrzynski et al., 2020). Additionally, arrays of unidentified proteins were observed in the outer matrix-facing PTT membranes, along with additional less organized protein densities. The arrays extend 5 nm globular domains into the PTT lumen, which based on comparison with AlphaFold models, is consistent with the overall size of CsCAH3. However, no membrane-binding domain is predicted in CsCAH3. Thus, if the globular domain is CsCAH3, it would have to be tethered to another protein in the membrane. If the arrays are CsCAH3, this would position CO_2_ release directly adjacent to the Rubisco matrix.

### Rubisco as a central organizing hub in diverse pyrenoids

PMTT, RHOP and PMST were identified through the presence of conserved RBM domains found in CsLinker. In the *Chlamydomonas* pyrenoid, Rubisco plays a central role as an organizing hub with multiple proteins containing EPYC1-type RBMs that have diverse structural functions (Meyer et al., 2020). It is still unclear is RBMs are primarily used for targeting proteins to the pyrenoid or for pyrenoid assembly and architecture. Although, the presence of RBM containing proteins with both catalytic and structural domains in both *Chlorella* and *Chlamydomonas* suggests a multipurpose role. It will be exciting to see if similar Rubisco-centric organizing principles (Wostrikoff and Mackinder, 2023; Ang et al., 2023) are found across diverse lineages including secondary red plastid algae such as diatoms.

The fusion of a starch binding CBM20 domain to a primarily disordered RBM containing region along with supporting localization and biochemical data, suggested that PMST may function analogously to SAGA1 in *Chlamydomonas* (Meyer et al., 2020; Crans et al., 2026). However, the lack of complementation in functional assays indicates that the mechanism of starch recruitment to the pyrenoid may differ between the two organisms, warranting further investigation. Moreover, recent studies have demonstrated that SAGA1 has a more complex multi-functional role where it operates alongside MITH1 to recruit membranes into the pyrenoid matrix (Hennacy et al., 2024).

The specific localization of PMTT to the PTT, combined with its high abundance, highlights its functional importance within the *Chlorella* pyrenoid. Given the presence of a transmembrane domain and multiple RBMs, we initially hypothesized that PMTT facilitates the recruitment of thylakoid membranes into the matrix. However, heterologous expression of PMTT failed to rescue the phenotype of the *Chlamydomonas mith1* knockout strain, which lacks traversing membranes. This inability to rescue the *mith1* phenotype suggests that instead of driving membrane recruitment, PMTT may function to stabilize the PTT around the matrix. Nevertheless, this lack of functional complementation in *Chlamydomonas* must be interpreted in light of the distinct structural differences between the pyrenoid membrane networks of these two green algae. We propose that PMTT is structurally specialized for the recruitment or stabilization of continuous thylakoid sheets, and it may lack the mechanistic capacity to facilitate the tubule formation and network recruitment required in *Chlamydomonas*.

### Rhodanese domain containing proteins are found in diverse pyrenoids

Both PMTT and RHOP contain rhodanese domains with the RHOP rhodanese domain predicted to be catalytically active and the PMTT domain inactive. Multiple rhodanese domain-containing proteins, including RBMP2, STR16 and STR18 have been identified in the *Chlamydomonas* pyrenoid (Lau et al., 2023). Rhodanese domains are known to interact with Fe–S cluster assembly proteins by supplying sulphur as a cofactor (Rydz, Wróbel and Jurkowska, 2021). Many Fe–S cluster assembly and Fe–S cluster containing proteins are reported to localize to the pyrenoid, where the hypothesized local oxygen limited conditions could facilitate their reactions (Lau et al., 2023). The conservation of catalytically active rhodanese proteins in the pyrenoid across lineages implies an important functional role. Recently it has been shown that the catalytically inactive rhodanese domain of RBMP2 is critical for extension of the pyrenoid tubules to form the reticulated region in *Chlamydomonas* (Wu et al., 2026). Whilst there is not an equivalent structure in *Chlorella*, it hints that the PMTT rhodanese domain may have a structural role in PTT formation.

### A structural and functional model of the *Chlorella* pyrenoid

Based on our cumulative observations, we propose a model of the pyrenoid-based CCM in the green microalga *Chlorella* (Fig. 6). Under CO₂ limiting conditions, starch granules form two, or in some cases three, plates that enclose the pyrenoid matrix. Cryo-ET further shows that two appressed thylakoid membranes (PTTs) traverse the pyrenoid matrix, dividing it into approximately two halves. Small fenestrations within the membranes provide direct connections between the pyrenoid matrix and the interthylakoid stroma outside of the pyrenoid (Fig. 2a,c), perhaps serving to channel ATP and Rubisco substrate and products between the stroma and pyrenoid, analogous to the proposed function of minitubules in *Chlamydomonas* (Engel et al., 2015).

**Figure 6:**
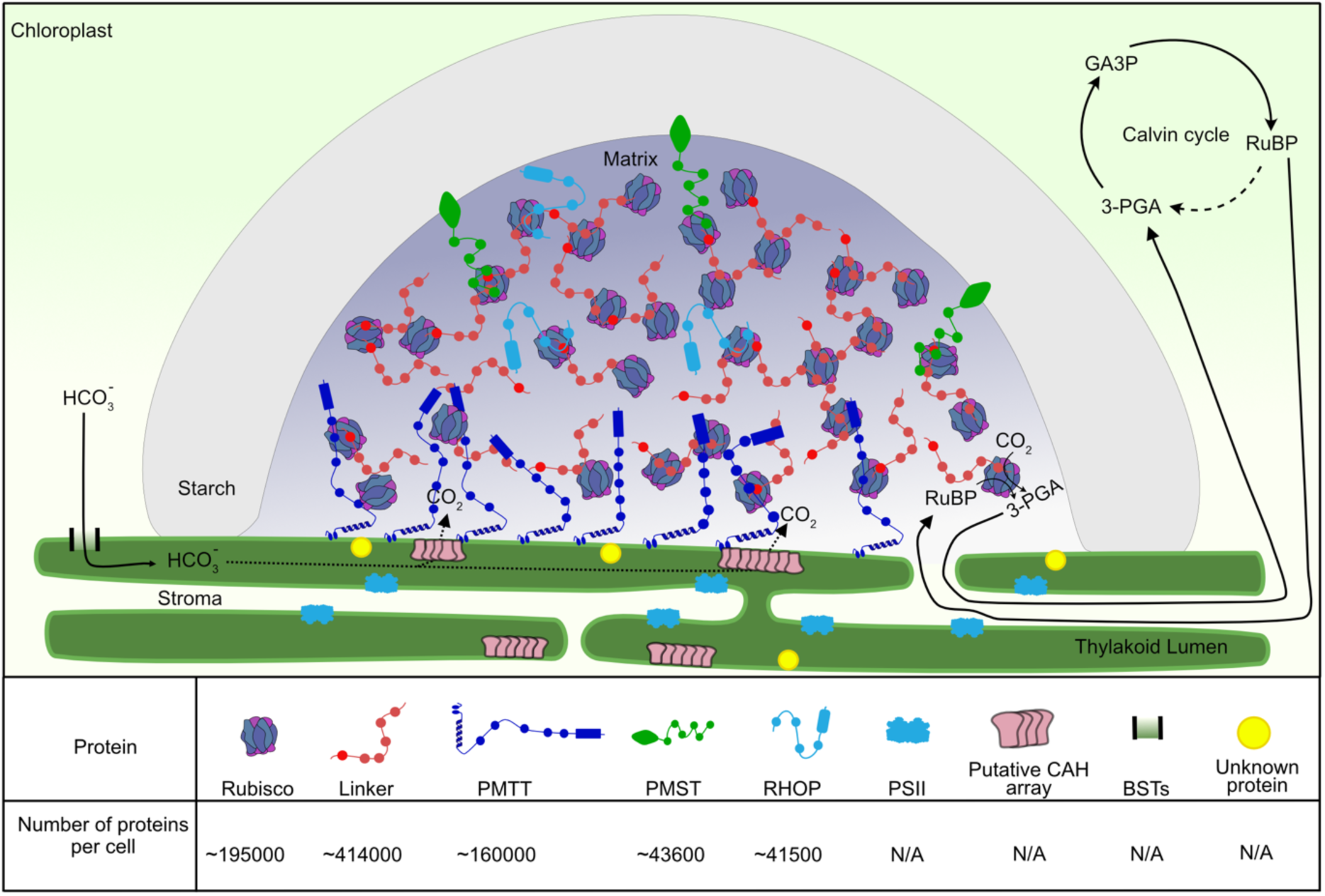
Proposed pyrenoid model in the green microalga *Chlorella sorokiniana* with protein abundances per cell from quantitative mass spectrometry shown where determined.

In *Chorella*, PTT membranes facing the stromal gap between appressed thylakoids harbour PSII complexes, whereas outer matrix-facing membranes contain ordered protein arrays, potentially composed of carbonic anhydrase CsCAH3 to accelerate the conversion of HCO_3_^-^to CO_2_ in close proximity to tightly packaged Rubisco in the pyrenoid matrix. In addition to these assemblies, smaller unassigned densities are also observed on these outer membranes. To link our cryo-ET observations to pyrenoid function, definitive identification of these membrane-associated proteins and clarification of the functional significance of PSII will be required.

With regard to protein components of the pyrenoid, CsLinker is highly abundant, found at an approximate 2:1 molar ratio with the Rubisco holoenzyme. Previous work has demonstrated that it is a functional analog of EPYC1 and mediates condensation of Rubisco to form the *Chlorella* pyrenoid (Barrett et al., 2024). PMTT’s interaction with liposomes composed of native lipids, its immunofluorescence localization to isolated pyrenoids, its localization to the central knot region of the *Chlamydomonas* pyrenoid and comparable abundance to the Rubisco holoenzyme, suggests a key functional role in initiating or stabilizing thylakoid membrane traversions through the matrix. PMST appears to be involved in recruiting starch to the periphery of the matrix formed by CsLinker–Rubisco condensates. This is supported by its ability to bind starch and facilitate the colocation of condensates onto starch plates. This structural architecture enables the delivery of HCO_3_^-^ by conserved BSTs into the thylakoid lumen where the enrichment of CsCAH3 in the PTTs ensures CO_2_ delivery to Rubisco within the matrix, with CO_2_ leakage out of the pyrenoid minimized by the starch sheath.

### *Chlorella* provides an expanded toolbox for plant pyrenoid engineering

Efforts to engineer a pyrenoid into plants to enhance productivity have, until now, predominantly relied on pyrenoid components identified from *Chlamydomonas* (Atkinson et al., 2016, 2020; Hennacy et al., 2024). A major limitation to this approach has been the lack of knowledge regarding pyrenoid structural components outside of *Chlamydomonas*. In this study, we characterize the pyrenoid from *Chlorella*, identified multiple potential pyrenoid proteins and characterized three of them, expanding the molecular toolbox for plant CCM engineering efforts. Expression analysis of these proteins in *N. benthamiana* demonstrated their expected localization, representing a critical step toward developing an active pyrenoid in plants. Future studies achieving stable expression of these proteins in plant cells harbouring a proto-pyrenoid will be critical to assess their functional contributions. For example, PMST may facilitate the recruitment of starch plates around the proto-pyrenoid, while PMTT could play a role in recruiting or stabilizing membranes within the matrix. Such experiments will be essential to evaluate the potential of these proteins for engineering a fully functional pyrenoid-based CCM in plants, either from *Chlorella* components alone or by creating a chimeric pyrenoid that combines convergently-evolved parts from diverse lineages such as *Chlorella*, *Chlamydomonas*, hornworts and diatoms.

## Acknowledgements

The University of York Department of Biology Technology Facility is acknowledged for providing access/support for fluorescence microscopy and proteomics. All current and past members of the Mackinder lab are thanked for fruitful discussions. Funding to L.C.M.M was from a UKRI Future Leaders Fellowship (MR/T020679/1 + MR/Y034074/1) and EPSRC (EP/W024063/1). Funding to L.C.M.M and A.J.M. was from a Carbon Technology Research Foundation (AP23-1_023), BBSRC-NSF/Bio (BB/Y000323/1) and Bill & Melinda Gates Foundation and United Kingdom Foreign, Commonwealth and Development Office (INV-054558). Funding to L.C.M.M and B.D.E from a BBSRC/Swiss National Science Foundation (SNSF) Partnering award (BB/X004953/1). Funding to A.J.M was from BBSRC (BB/W003538/1, BB/Y008162/1). Funding to M.I.S.N was from the NWO (Dutch Research Council) under the Rubicon Postdoctoral Fellowship from December 2022-November 2024 (Project ID: 019.221EN.010). P.V.d.S, M.H., and B.D.E. were funded by Swiss Nanoscience Institute PhD School grant P2204. Additional funding to B.D.E. was from ERC consolidator grant “cryOcean” (fulfilled by the Swiss State Secretariat for Education, Research, and Innovation, M822.00045).

The authors would like to acknowledge Dr. Ousmane Dao for his technical assistance and support in conducting the oxygen evolution experiments and Dr. Philipp Girr for his work which established the foundation for the liposome–condensate interaction studies presented in this article. We would also like to thank all current and past Mackinder Lab members for their discussions and input throughout the project. At the University of Basel, cryo-ET sample preparation and acquisition were performed at the BioEM facility, and cryo-ET computational analysis used the sciCORE scientific computing center (http://scicore.unibas.ch/).

## Author Contribution

M.I.S.N, J.B and L.C.M.M conceived the study and wrote the manuscript, with contributions from P.V.D.S, M.D, J.P, B.D.E and A.J.M. M.I.S.N performed oxygen evolution experiments, TEM, Mass Spectrometry, absolute quantification, biochemical characterization of proteins and in vivo *Chlamydomonas* experiments. A.D supported with the absolute quantification mass spectrometry experiments. J.B identified the candidate proteins, performed pyrenoid extraction and immunofluorescence for *Chlorella*. P.V.D.S and M.D performed the Cryo-ET and processed the data with input from B.D.E. J.P performed all the *in planta* expression of proteins with input from A.J.M. T.G developed the methods for liposome preparation.

## Data availability

Proteomics data for VLC and HC grown cells and absolute protein quantification will be deposited in MassIVE, and given ProteomeXchange identifiers.

Rubisco co-IP data was reanalyzed from (Barrett et al. 2024), with ProteomeXchange identifier PXD044179.

Raw cryo-ET data + reconstructed/denoised tomograms will be deposited in EMPIAR and EMD.

Subtomograms averages will be submitted on EMDB.

## Methods

### Strains and culture conditions

*Chlamydomonas reinhardtii* (CC125), *Chlamydomonas reinhardtii* (CC4533), ΔMITH1, ΔEPYC1, ΔSAGA1, *Chlorella sorokiniana* UTEX1230 (SAG 211-8K) and all resulting strains were maintained on Tris-acetate-phosphate media plates constituted with 1.5% Agar and revised trace elements at low light conditions (∼10 μmol photons m^−2^ s^−1^) at 21°C. For TEM and OER experiments the cultures were grown in HC conditions and then adapted for 24 hours in LC or VLC in Tris-Phosphate (TP) media before harvesting and using for experiments. For comparative proteomic mass spectrometry, *Chlorella* cells were cultured in 500 mL bottles containing 300 mL of TP media with continuous sparging of HC or VLC gases. Cells were harvested during the exponential growth phase by centrifugation at a density of 10^7^ cells per growth condition. The harvested cell pellets were washed with phosphate-buffered saline (PBS) and subsequently resuspended in 200 µL of lysis buffer consisting of 5% SDS and 50 mM TEAB (pH 8.5). Cell lysis was achieved via sonication at 10A for three 15-second intervals, with samples kept on ice for more than 1 minute between cycles to cool. The resulting cell lysate was centrifuged at 20,000 g for 20 minutes, after which the supernatant was collected. Finally, the supernatant was frozen at -80°C prior to mass spectrometry submission

### Photosynthetic Kinetics Measurement Using Oxygen Evolution

The K_0.5_ and V_max_ for *Chlorella* and *Chlamydomonas* (strain CC-125) were determined from oxygen evolution rate measurements using a Hansatech OxyLab+ system. Cultures were acclimated to either HC, LC or VLC conditions in TP (pH 7.4) and harvested during the exponential growth phase by centrifugation at 1800 g for 3 minutes. Chlorophyll content was quantified, and culture volumes were adjusted to have 50 µg Chl mL^−1^ in 2 mL of fresh medium sparged with CO_2_ free air.

Resuspended cells were transferred to a thermostatically controlled reaction chamber maintained at 22 °C and equipped with an Oxygraph+ oxygen electrode system (Hansatech, Norfolk, England). To eliminate background oxygen evolution, cultures were initially sparged with CO₂-free air. Photosynthetic activity was then stimulated by exposing the culture to 750 µmol photons m^−2^ s^−1^ and sequential addition of sodium bicarbonate to achieve final concentrations of 1, 2, 3, 5, 10, 25, 50, 100, 250, and 500 µM. Oxygen evolution was continuously recorded after each bicarbonate addition, allowing for the generation of photosynthetic response curves and calculation of kinetic parameters.

### Antibodies specific for candidate proteins

Polyclonal antibodies (synthesised by YenZym Antibodies) specific to the candidate pyrenoid proteins were raised in rabbits using peptides corresponding to unique antigenic regions of each protein. These antibodies were subsequently used in immunogold labeling, immunofluorescence microscopy, co-immunoprecipitation and protein lipid overlay assays. The peptide sequences used for immunization were as follows: RbcL, EVWKEIKFEFETIDTL-COOH; CsLinker, PTPVSNSGVRSAMSSG-amide; PMTT, SYSPPPPSQQQAWEPAQ-amide; RHOP, TPAPAAPSYSYGSAS-amide; and PMST, SSPRPPAVRRSSS-amide. As a negative control, an antibody raised against the *Chlamydomonas* protein BST2 using the peptide PDLDSINAAAPNGNGSHNGN-amide was used.

### Transmission electron microscopy and Immunogold assay

Cells cultured in TP medium under HC, LC and VLC conditions were harvested by centrifugation and immediately fixed in 2.5% (v/v) glutaraldehyde prepared in TP medium by nutating for 1 hour at room temperature. Following fixation, the cells were pelleted and osmicated with 1 mL of a freshly prepared solution containing 1% osmium tetroxide (OsO₄), 1.5% (w/v) potassium ferricyanide (K₃Fe(CN)₆), and 2 mM calcium chloride (CaCl₂), for 2 hours with continuous rotation at room temperature. After osmication, the cells were washed thoroughly with distilled water (4–5 washes) and subjected to a graded ethanol dehydration series by incubating sequentially for 5 minutes each in 30%, 50%, 70%, and 95% ethanol. Final dehydration was performed using 100% ethanol for 10 minutes, followed by two washes in 100% acetonitrile for 10 minutes each. The dehydrated pellets were embedded in 1:3, 1:1 and 3:1 Spurr resin to acetonitrile without catalyst overnight, followed by incubation in 100% Spurr resin with catalyst for 24 hours. The resin was refreshed daily with fresh catalysed Spurr resin for four consecutive days. Polymerisation was performed at 70 °C for 24 hours. The resulting blocks were sectioned using a Leica UCT7 ultramicrotome with a Diatome diamond knife, and ultrathin sections were mounted on nickel grids. Imaging was performed using a Thermo Fisher Talos L120 G2 TEM using an accelerating voltage of 120kV and micrographs were obtained using a bottom-mounted Thermo Fisher Ceta 16 megapixel TEM camera.

For immunogold labelling, following glutaraldehyde fixation, specimens were directly dehydrated through a graded ethanol series (10% to 70% ethanol), allowing 15 minutes between each step. From 30% ethanol onward, dehydrations were conducted at −10 °C. Subsequent resin infiltration was also carried out at −10 °C using graded mixtures of LR White resin and 70% ethanol (ratios of 1:3, 1:1, and 3:1), with 1 hour of incubation per step. A final overnight infiltration in 100% LR White resin was performed. Specimens were then transferred to fresh LR White resin in gelatine capsules and polymerised under UV light at −10 °C for 48 hours. The ultrathin sectioned and grid-mounted samples were given 1 hour at room temperature exposed to 1% sodium metaperiodate aq., rinsed with double distilled water and then blocked with 3% bovine serum albumin (BSA) in PBS for 30 minutes at room temperature. Following blocking, the grids were incubated with the primary antibody (1:5000 dilution) for 1 hour, and then incubated for an additional 1 hour with a goat anti-rabbit IgG secondary antibody conjugated to 10 nm gold particles (Merck). After final washes, the samples were prepared for transmission electron microscopy imaging by 5 minutes of staining with sequential solutions of 2% uranyl acetate in 30% ethanol/aq. and Reynolds lead citrate (protected by sodium hydroxide pellets during this step). Staining solutions were thoroughly rinsed from grids using degassed double distilled water.

### Cryo-ET sample preparation and data acquisition

A total of 4.5 µL cell suspension (1500–2000 cells/µL), grown in TAP media at air level CO_2_ and 50 µE light, were applied to a holey carbon R2/1 grid (Quantifoil). Cells were plunged frozen in liquid ethane with back-sided blotting using a Leica EM GP2 (2 s blot time and 90% humidity). Samples were stored under liquid nitrogen conditions.

FIB milling was performed as described in (Schaffer et al., 2017). Briefly, grids clipped in Autogrids (Thermo Fisher Scientific) were loaded into an Aquilos 2 microscope (Thermo Fisher Scientific) and a layer of organometallic platinum was applied using the gas injection system. For milling and imaging, the gallium ion beam was operated at 30 kV.

Tilt-series were acquired on a Titan Krios G4 transmission electron microscope operating at 300 kV equipped with a Selectris X energy filter (slit set to 10 eV) and a Falcon 4i direct electron detector (Thermo Fisher Scientific), recording dose-fractionated TIFF movies. A dose-symmetric tilt scheme was used in a tilt span of ± 54°, covered by 2° steps starting at either ± 10° offset to compensate for the lamella pre-til set by the TEM Tomography 5 software (Thermo Fisher Scientific). The 100 μm objective aperture was inserted during collection. Target defocus was set for each tilt-series in a range of -2 µm to -4 µm in steps of 0.5 µm. The microscope was set to a magnification of either 67’000x or 53’000x corresponding to a pixel size of 1.91 Å or 2.42 Å respectively at the sample level and a nominal dose of 2 e-/Å2 per tilt image.

### Cryo-ET data processing

Tomograms were processed with Scipion (Jiménez de la Morena et al., 2022) using the workflow described in (Github). Briefly, raw frames were motion corrected using Motioncorr3 (Zheng et al., 2017), and images were aligned after manual tilt-series curation with either AreTomo2 or IMOD’s etomo (Kremer, Mastronarde and McIntosh, 1996). CTF-corrected (by phase-flipping), dose filtered, and four times binned tomograms were reconstructed using IMOD (Kremer, Mastronarde and McIntosh, 1996).

Tomograms were denoised from their reconstructed half-volumes using DeepDeWedge (Wiedemann and Heckel, 2024) and membrane segmentation was done with Membrain-seg (Lamm et al., 2024). Flat membranes were created and visualized using Surforama (Yamauchi et al., 2024). Rubisco positions were mapped back from template matching CTF-corrected tomograms using pytom-match-pick (Chaillet et al., 2025). Starch was masked manually in Amira (Thermo Fisher Scientific).

Initial putative CAH3 positions were manually picked in ArtiaX (Ermel, Arghittu and Frangakis, 2022) normal to the membrane and exported as RELION5 (Burt et al., 2024) particles. Tomogram alignments were imported using a custom python script (Van der Stappen). An initial reconstruction was used as a reference for subsequent rounds of 3D refinement using in-plane searches (--sigma_tilt 3 --sigma_psi 3) and 3D classification (4 classes, T=0.2, 25 iterations). A total of 896 particles from 9 tomograms was used for CTF refinement and Bayesian polishing which yielded a final postprocess map at 20 Å based on gold-standard FSC.

### Bioinformatic identification of *Chlorella* pyrenoid proteins

To identify potential pyrenoid proteins in *Chlorella*, the Rubisco-binding region (RBM) of CsLinker (APAASYTAPAAAPAGGLSAEQREFLERKARESRSPAPVETRGRSAPR) was used as a query to BLAST against the *Chlorella* proteome. This search identified two candidate proteins, PMTT and RHOP.

To identify PMST, a potential homolog of SAGA1, the carbohydrate-binding module 20 (CBM20) domain of SAGA1 (E = 0.01; SAFTLCRFVVPSDPPAPLPHPSSSTAAAAAAATDVLFLVGSCPELGEWDPGRALKLAAVAGGGWAAEARLELESEVAAKLLIMRDGTRMEWELGPNRVLRGALTAAAAQPGTGAPPPRALLFSCPWN) was used as a BLAST query against the *Chlorella* proteome. This search returned nine proteins. Among these, one protein (CSI2_123000007310) contained an N-terminal CBM20 domain and RBMs with sequence homology to CsLinker.

### Pyrenoid isolation mass Spectrometry

Cells were grown to mid-exponential phase (2x10^6^ cells ml-^1^) in TP media under ambient CO_2_ levels. Cultures were harvested by centrifugation (1500g, 10 min) and resuspended in 30 mM HEPES-KOH (pH 8.0). Following a second pelleting step, cells were fixed in 1 mL of 30 mM HEPES-KOH (pH 8.0) containing 1%(w/v) formaldehyde for 20 minutes under gentle rotational mixing. The fixation was quenched by the addition of 1 mL of 2M Tris-HCl (pH 8.0). The fixed cells were washed twice and resuspended in pyrenoid enrichment buffer (50 mM Tris-HCl, 0.2 mM EDTA, 0.5%[w/v] Triton X-100, pH 8.0) to a final volume of 1 mL. 1 mL of the partially fixed cells are lysed by sonication using a Misonix-4000 sonicator and 1/16” diameter sonication probe operated at 30% amplitude with setting 3 second ON and 6 seconds OFF for a total processing time of 3 minutes. The pyrenoid is enriched from the lysed sample by centrifugation at 2500g for 20 minutes at room temperature. Wash the pelleted pyrenoid enriched fraction in 1 mL of pyrenoid enrichment buffer and resuspend to 1 mL in pyrenoid enrichment buffer. Load the crude pyrenoid fraction onto a 9 mL cushion of 100% Percoll in a 15 mL falcon tube and centrifuge at 2500 g for 15 minutes at room temperature to pellet the pyrenoid-enriched fraction. The pyrenoid fraction was washed once in the pyrenoid enrichment buffer and finally resuspended in a buffer containing 50 mM Tris-HCl (pH8.0) and 50mM NaCl. The resulting enriched pyrenoid fractions were then processed for mass spectrometry (MS) analysis.

### Immunofluorescence of pyrenoids

Pelleted pyrenoid-enriched fractions were resuspended in 100 µL of 1% (w/v) bovine serum albumin (BSA) in Tris-buffered saline with Tween (TBST) containing the primary antibody in 1:50 dilutions. The pyrenoid suspension was incubated overnight at 4°C with gentle mixing and washed twice with 1 mL of TBST. Samples were then resuspended in 100 µL of 1% (w/v) BSA in TBST containing a 1:1000 dilution of fluorescent secondary antibody (anti-rabbit A-11008 or anti-mouse T6074, Sigma-Aldrich) and incubated for 1 hour at 4°C. After incubation, the fractions were washed twice with TBST and subsequently imaged using fluorescence microscopy.

### Co-immunoprecipitation mass spectrometry (Co-IP/MS)

Co-IP/MS was replotted from (Barrett et al., 2024). The BST2 antibody was used as a control.

### Protein purifications

Rubisco purification from *Chlorella* biomass was carried out using cultures grown to the late exponential phase in 10 L of TP medium sparged with high CO₂ (HC). The harvested biomass was resuspended in lysis buffer containing 100 mM NaCl, 100 mM Tris-HCl (pH 8.0), 0.5 mM EDTA, 1 mM DTT, 2× protease inhibitor tablets (Roche), and 2 mM PMSF. Cell lysis was performed by ultrasonication for 10 minutes using a cycle of 3 seconds ON and 10 seconds OFF. The lysate was cleared by centrifugation at 30,000 × g for 30 minutes using a Lynx high-speed centrifuge to remove cell debris. The resulting supernatant was used for Rubisco purification, which was carried out as previously described by Barrett et al. (2024).

The pyrenoid proteins CsLinker, PMTT, PMST, and RHOP were heterologously expressed in *E.coli* BL21 cells with MBP and GFP solubility tags and purified using an ÄKTA FPLC system. Overnight pre-cultures grown in LB medium were diluted 1:100 into 1 L of fresh LB and incubated at 37 °C until reaching an optical density at 600 nm (OD₆₀₀) of ∼0.5. Protein expression was induced with IPTG, and cultures were incubated for an additional 4 hours at 37 °C. Cells were harvested by centrifugation and resuspended in 15 mL of lysis buffer containing 50 mM Tris-HCl (pH 8.0), 500 mM NaCl, 25 mM imidazole, and 2× protease inhibitor tablets (Roche). Cell lysis was performed by ultrasonication (7 min total processing time, 3 s ON/12 s OFF cycle). Lysates were cleared by centrifugation at 50,000 × g for 30 minutes using a Lynx high-speed centrifuge. The resulting supernatant was passed through a 50 µm filter to remove particulates before chromatography.

Protein purification was initiated with immobilised metal affinity chromatography (IMAC) using a 5 mL HisTrap column (Cytiva). Elution was carried out using a 100 mL (20 column volumes) linear gradient from lysis buffer (without protease inhibitors) to elution buffer containing 50 mM Tris-HCl (pH 8.0), 500 mM NaCl, and 500 mM imidazole. Eluted target proteins were incubated with TEV protease overnight at 4 °C to cleave the solubility tags (MBP and GFP). The sample was subsequently buffer-exchanged to remove imidazole and re-applied to the HisTrap column to separate the cleaved tags and TEV protease from the protein of interest. The flow-through, containing the untagged protein, was collected and further purified by size exclusion chromatography using a HiLoad 16/600 Superdex 75 pg column (Cytiva).

For PMTT, the lysis buffer was supplemented with 6 M urea to solubilise the protein, as denaturing conditions were necessary for its purification.

### Absolute quantification mass spectrometry

Absolute quantification of *Chlorella* pyrenoid proteins was performed following the method reported by Barrett et al. (2024). Purified proteins: PMTT, PMST, and RHOP; were used to generate standard curves for quantifying their respective abundances in *Chlorella* cells. Rubisco and CsLinker data are from Barrett et al. (2024).

### Droplet sedimentation assay

Sedimentation assays were performed in a buffer containing 50 mM Tris-HCl (pH 8.0) and 50 mM NaCl. Each reaction was carried out in a total volume of 5 µL in PCR tubes. After protein addition, samples were incubated at room temperature for 15 minutes, followed by centrifugation at 10,000 × g for 10 minutes. The supernatant was carefully transferred to a fresh tube for analysis of the soluble fraction. To the original tube containing the pellet, 1x Laemmli loading buffer was added to solubilize the pellet fraction. Both soluble and pellet fractions were analyzed by SDS–PAGE.

### In vitro confocal microscopy

All in vitro confocal imaging was performed using a Zeiss LSM880 confocal microscope equipped with a 63×/1.4 NA Plan-Apochromat oil immersion objective (Zeiss). Image acquisition and system operation were conducted using Zen Blue software (Zeiss). Samples were mounted in µ-Slide 15-well 3D chambers (Ibidi) for imaging.

### Starch purification from *Chlorella*

The method for starch extraction was adapted from Delrue et al. (1992). A 50 mL culture of *Chlorella* cells in the late exponential growth phase (∼5 × 10⁸ cells), grown in TP medium, was harvested by centrifugation at 3,000 × g for 10 minutes. The cell pellet was resuspended in 15 mL of starch extraction buffer containing 50 mM Tris-HCl (pH 8.0), 0.2 mM EDTA, and 0.5% (v/v) Triton X-100. The suspension was then sonicated using a Misonix ultrasonic liquid processor equipped with a microtip (suitable for 1–15 mL volumes) for a total of 10 minutes, applying a 30-second ON / 30-second OFF cycle at 25 Amp.

The lysate was centrifuged at 3,000 × g for 10 minutes, and the resulting pellet was resuspended in 5 mL of extraction buffer. This 5 mL suspension was layered on top of 5 mL of a 95% Percoll cushion in a 15 mL Falcon tube and centrifuged again at 3,000 × g for 10 minutes. The starch pellet, visible as a white deposit at the bottom, was collected and resuspended in 5 mL of extraction buffer. The Percoll step was repeated once to increase purity. Finally, the pellet was washed twice with 2–5 mL of distilled water and used for downstream assays.

### Starch-Protein(Condensate) colocalization

Z-stack images of starch and protein/condensate colocalization were analyzed in Fiji. For analysis, the slice providing the clearest view of starch granules was selected. Image channels were split to separate the fluorophore signal from the starch visible in the bright-field channel. The starch channel was processed using the Otsu auto-thresholding method to extract starch signals from the bright field. Binary image processing steps (Fill Holes, Watershed, and Close) were applied to refine the starch selection. Using the Analyze Particles function, starch granules ranging from 1 to 100 µm in size were identified, as larger structures corresponded to multiple granules clumped together. These selections were saved as regions of interest (ROIs) in the starch channel. The ROIs were then overlaid onto the fluorophore channel, and fluorescence intensity within each ROI was measured to assess colocalization. Background fluorescence was estimated by measuring intensity in a manually selected ROI placed outside regions containing fluorescence and subtracted to obtain corrected intensity values. Starch granules with corrected fluorescence intensities positive for colocalization with protein/condensate signals are quantified, as shown in Figure 4.

### Protein-Lipid Overlay (PLO) assay

Membrane lipid strips (P-6002; Echelon Biosciences) were used to assess protein–lipid interactions. The strips were first blocked for 1 hour at room temperature in a TBST buffer containing 3% (w/v) bovine serum albumin (BSA). Following the blocking step, ∼100 µg of purified protein was diluted in fresh TBST-3% BSA solution and incubated with the lipid strips for 2–3 hours at room temperature with gentle shaking.

After incubation, the strips were washed twice for 10 minutes each in TBST-3% BSA buffer with gentle agitation. The membranes were then incubated with the protein-specific primary antibody (1:1000 dilution in TBST-3% BSA) for 1 hour at room temperature. After three subsequent washes (10 minutes each) in TBST-3% BSA, the strips were incubated for 1 hour with a secondary antibody (Goat anti-Rabbit IgG (H+L), Cross-Adsorbed, Alexa Fluor® 488 from Thermo Fisher Scientific) diluted 1:10,000 in TBST-3% BSA.

Finally, the strips were washed twice in TBST-3% BSA and imaged using an Amersham Typhoon Biomolecular Imager with Cyt2 settings.

### Liposome preparation

All lipids used in this study were purchased from Avanti Polar Lipids: 18:2 monogalactosyldilinoleoylglycerol (MGDG), digalactosyldiacylglycerol (DGDG), sulfoquinovosyldiacylglycerol (SQDG), 1,2-dioleoyl-sn-glycero-3-phospho-(1’-rac-glycerol) (DOPG), and 1,2-dioleoyl-sn-glycero-3-phosphoethanolamine-N-(lissamine rhodamine B sulfonyl) (LissRhdPE; ammonium salt).

GUV liposomes were used for in vitro confocal microscopy to investigate the interaction of proteins and condensates with liposomes. GUVs were prepared by optimizing previously reported protocols. A 30 µL aliquot of 5% (w/v) polyvinyl alcohol (PVA, fully hydrolyzed, Sigma Aldrich) solution was applied to a clean glass slide and spread evenly using a coverslip to form a uniform PVA film. The slide was then dried on a heat block at 60 °C for 1 hour. After cooling to room temperature, 10 µL of lipid mixture at 10mM concentration, prepared in chloroform was deposited onto the PVA-coated slide and gently spread with a coverslip to achieve a uniform lipid layer. The lipid-coated slide was dried under a nitrogen stream for 5 minutes, followed by further desiccation under vacuum for 2 hours.

For hydration, 300 µL of buffer containing 50 mM Tris-HCl (pH 8.0) and 50 mM NaCl was carefully added onto the dried lipid layer and incubated at room temperature, protected from light, for 1 hour to allow vesicle formation. The resulting suspension containing GUVs was gently transferred to a clean microcentrifuge tube and used immediately for downstream experiments.

### Plasmid cloning

For protein expression in Escherichia coli, the coding sequences of PMTT, PMST, and RHOP were codon-optimised using the TWIST codon optimisation tool (TWIST Biosciences). A previously developed plasmid, His-mEGFP-TEV-cloning site-TEV-MBP-His, was submitted to TWIST Biosciences to serve as the reference vector for cloning. The codon-optimised sequences were synthesised and inserted between the TEV sites by TWIST, resulting in the following expression constructs: His-mEGFP-TEV-PMTT-TEV-MBP-His, His-mEGFP-TEV-PMST-TEV-MBP-His, and His-mEGFP-TEV-RHOP-TEV-MBP-His. The construct His-mEGFP-TEV-PMTT-TEV-MBP-His was further modified to include the N-terminal amino acids that were initially omitted. The corresponding N-terminal coding region was codon-optimised for *Escherichia coli*, synthesised by TWIST Biosciences, and cloned in-frame at the 5’ end of the previously obtained PMTT sequence. The full-length PMTT coding sequence was subsequently inserted into the His-mEGFP-TEV-cloning site-TEV-MBP-His vector using Golden Gate assembly.

For chloroplast expression in *Chlamydomonas*, the pME_Cp_2_098 plasmid (a gift from René Inckemann and Tobias Erb) was used as the backbone vector. The *E. coli* codon-optimised coding sequences of PMTT, PMST, and RHOP were cloned into this plasmid using fluorescent reporters (mVenus or mCherry) as described in the plastome engineering toolbox (Inckemann et al., 2024).

For development of a co-expression construct in *Chlamydomonas* chloroplast, the same plastome toolbox based on Golden Gate cloning was employed. In the first round of Level 1 assembly, CsLinker (Barrett et al., 2024) and mCerulean (Inckemann et al., 2024) were PCR-amplified and cloned into pME_Cp_1_473_CDS-PH_GOI1/3_rbcL. Similarly, full-length PMTT and mVenus were cloned into pME_Cp_1_474_CDS-PH_GOI2/3_rbcL, and PMST and mScarlet were cloned into pME_Cp_1_475_CDS-PH_GOI3/3_rbcL. All three Level 1 constructs were assembled with pME_Cp_1_472_tobra-resistance_cassette_inverse_rbcl-locus and pME_Cp_0_7-8_005_Kan/ColE1_v2_mScarlet in a five-part Golden Gate reaction to generate the final Level 2 plasmid. This construct enabled simultaneous expression of CsLinker-mCerulean, PMTT-mVenus, and PMST-mScarlet in the *Chlamydomonas* chloroplast.

### *Chlamydomonas* chloroplast transformation

*Chlamydomonas* cells (1 × 10⁷) in the exponential growth phase were plated on TAP agar plates supplemented with 100 µg mL^-1^ hygromycin for selection. Chloroplast transformation was carried out using particle bombardment with gold nanoparticles. For each transformation, 1 µg of plasmid DNA was mixed with 0.5 mg of 550 nm gold particles following the manufacturer’s instructions. A Biolistic PDS-1000/He particle delivery system (Bio-Rad) equipped with a 1100 psi rupture disk was used to deliver the DNA-coated gold particles into the cells. Transformed colonies typically appeared within 5–7 days post-transformation.

Selected colonies were re-streaked onto fresh TAP plates containing 250 µg mL^-1^ hygromycin. Re-streaking was performed three times to enrich homoplasmic lines. Colonies from the final round were used for confocal microscopy, after growth in either TAP or TP media to an appropriate cell density.

For co-expression constructs, selection was based on tobramycin resistance. Initial selection was performed on TAP plates containing 100 µg mL^-1^ tobramycin. Colonies were sequentially re-streaked on TAP plates with 150, 200, and finally 250 µg mL^-1^ tobramycin to ensure stable integration and expression of the transgenes.

### *Chlamydomonas* confocal microscopy

All images of protein localisation were captured using a Zeiss LSM880 confocal microscope equipped with Airyscan detection. Imaging was performed with a 63× oil-immersion Plan-Apochromat objective lens (NA 1.4, Carl Zeiss). Samples were mounted in 18-well µ-slide chambers (Ibidi), where 10 µL of sample was immobilised by overlaying with 30 µL of 1.5% low-melting-point agarose prepared in TP medium.

### Plant construct design and *N. benthamiana* transformation

Coding sequences of PMTT, RHOP and PMST were codon optimised for Arabidopsis and synthesised as DNA fragments (GeneArt, Thermo Fisher Scientific) for assembly into level 0 vectors using the Plant MoClo system (Engler et al., 2014). PMTT, RHOP and PMST were each assembled into a level 1 expression vector with the 35s cauliflower mosaic virus (CaMV) promoter, a C-terminal mEGFP tag and HSP+nos double terminator (Diamos and Mason, 2018). A truncated variant of PMST (trncPMST) lacking 50 residues from the N-terminus was generated by PCR and assembled into a level 1 expression vector with the cassava vein mosaic virus (CsVMV) promoter, the AtRbcS1A (AT1G67090) chloroplast transit peptide consisting of 80 residues of the N-terminus of AtRbcS1A (cTP1A80), a N-terminal mCherry tag with a 9-residue flexible linker sequence and the HSP+nos terminator. Overnight cultures of electrocompetent *Agrobacterium tumefaciens* GV3101 carrying level 1 plant expression vectors were grown in LB under antibiotic selection at 30⁰C. Cultures were spun down and resuspended in MMA media (10 mM MES pH 5.6, 10 mM MgCl_2_, 0.1 mM acetosyringone) to OD 600 = 0.8 and syringe infiltrated into fully expanded leaves of 4-week-old *N. benthamiana* plants and incubated for 3 days at room temperature on the laboratory benchtop.

### Plant Confocal laser scanning and starch staining

*N. benthamiana* leaves were imaged using a Leica SP8 confocal microscope as in (Atkinson et al., 2016). Starch granules were stained by floating small explants of leaf in fluorescein (CAS No.: 2321-07-05) at a final concentration of 0.1 mM. Images were processed using Fiji (https://fiji.sc/).

## Extended Data Figures

**Extended Data Fig. 1.**
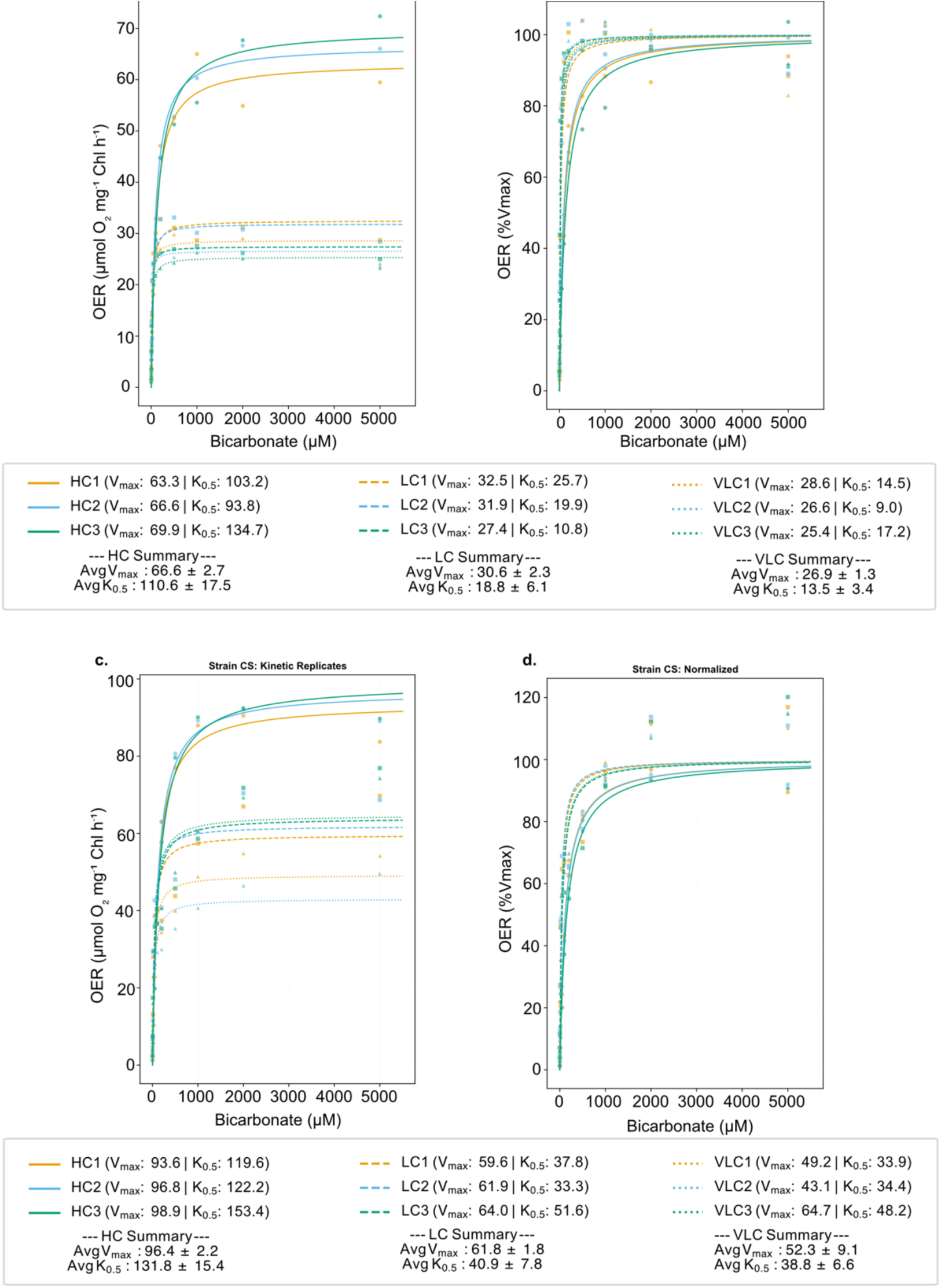
Net oxygen production in response to bicarbonate addition for a,b) *Chlamydomonas reinhardtii* CC125 (top) and c,d) *Chlorella sorokiniana* (bottom) adapted to high CO₂ (HC), low CO₂ (LC) and very low CO₂ (VLC). Normalized values for OER as %Vmax is shown on the right side for each culture. Cultures grown in TP medium were subjected to stepwise additions of bicarbonate, and oxygen evolution rate (OER) was measured over a 40-second interval immediately following each addition.

**Extended Data Fig. 2.**
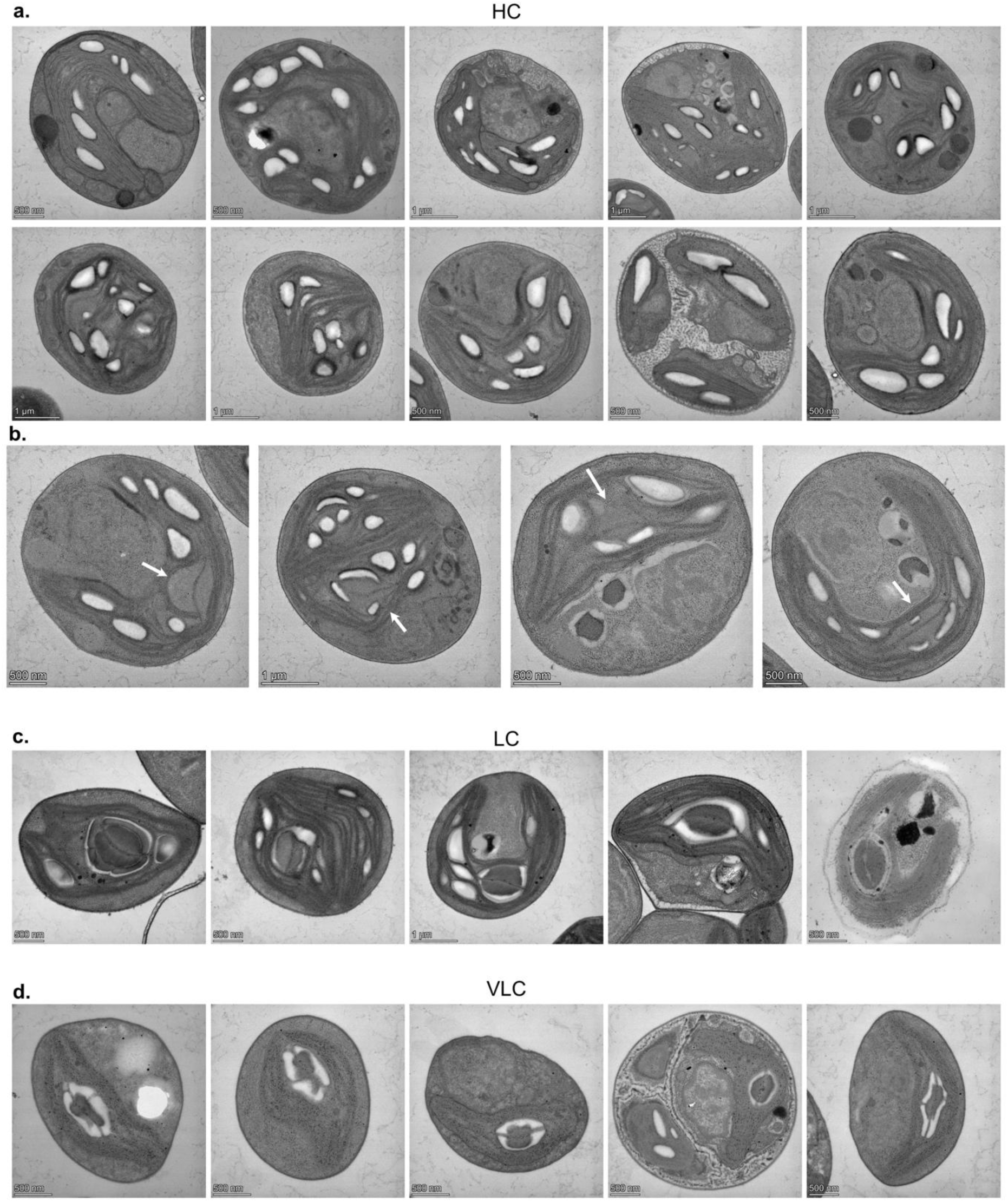
Transmission electron micrographs showing *Chlorella sorokiniana* cells adapted to different CO_2_ levels. a) High CO_2_ adapted cells with no visible pyrenoid. b) High CO_2_ adapted cells with a reduced pyrenoid containing a single thylakoid traversion (marked with white arrow). c) Low CO_2_ and d) Very low CO_2_ adapted cells.

**Extended Data Fig. 3.**
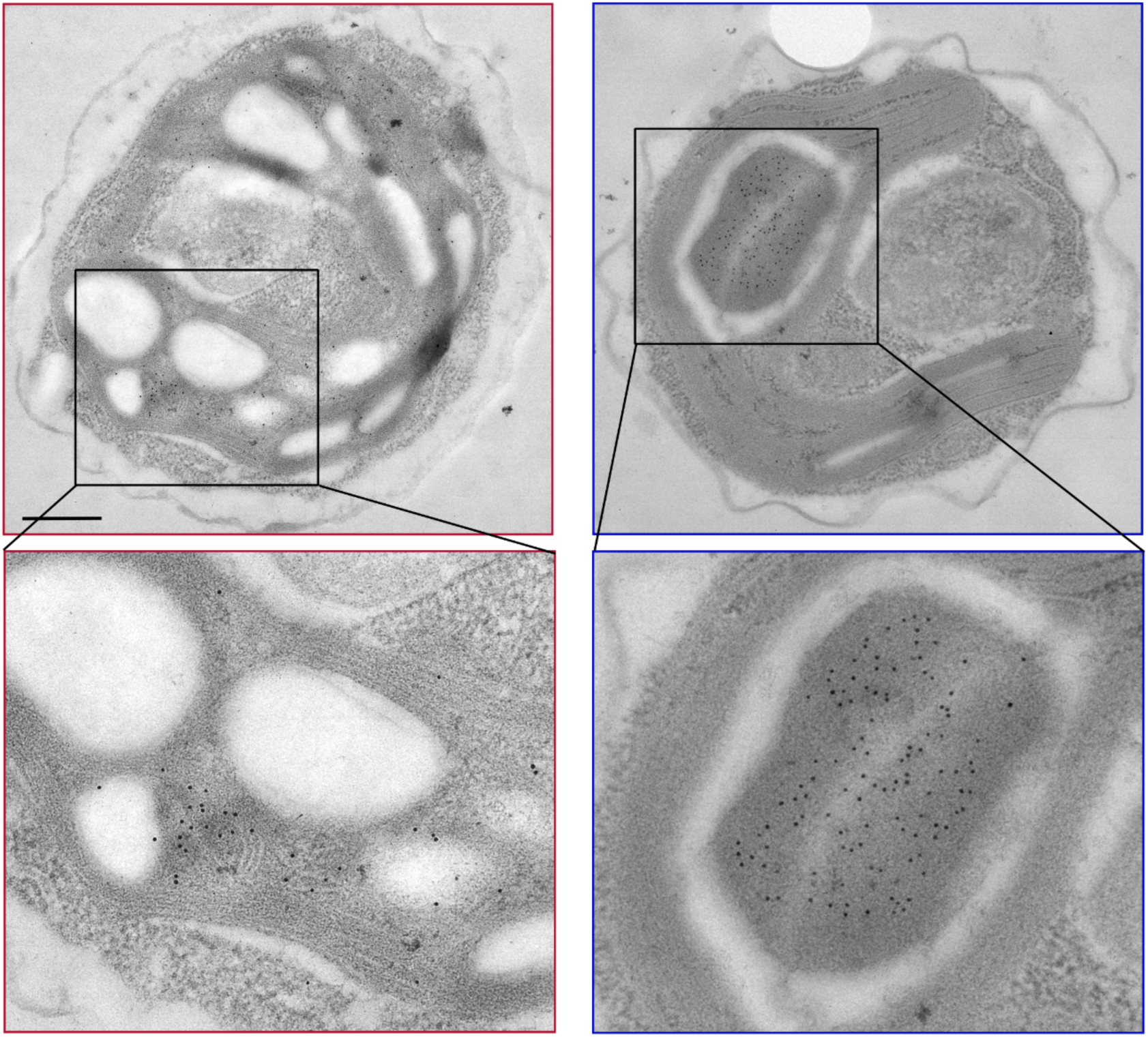
Transmission electron micrographs showing immunogold labelling of Rubisco large subunit in *Chlorella sorokiniana* cells adapted to high CO₂ (HC, left) and low CO₂ (LC, right) conditions. Top row are images used for Figure 1e. Scale bar: 500 nm.

**Extended Data Fig. 4.**
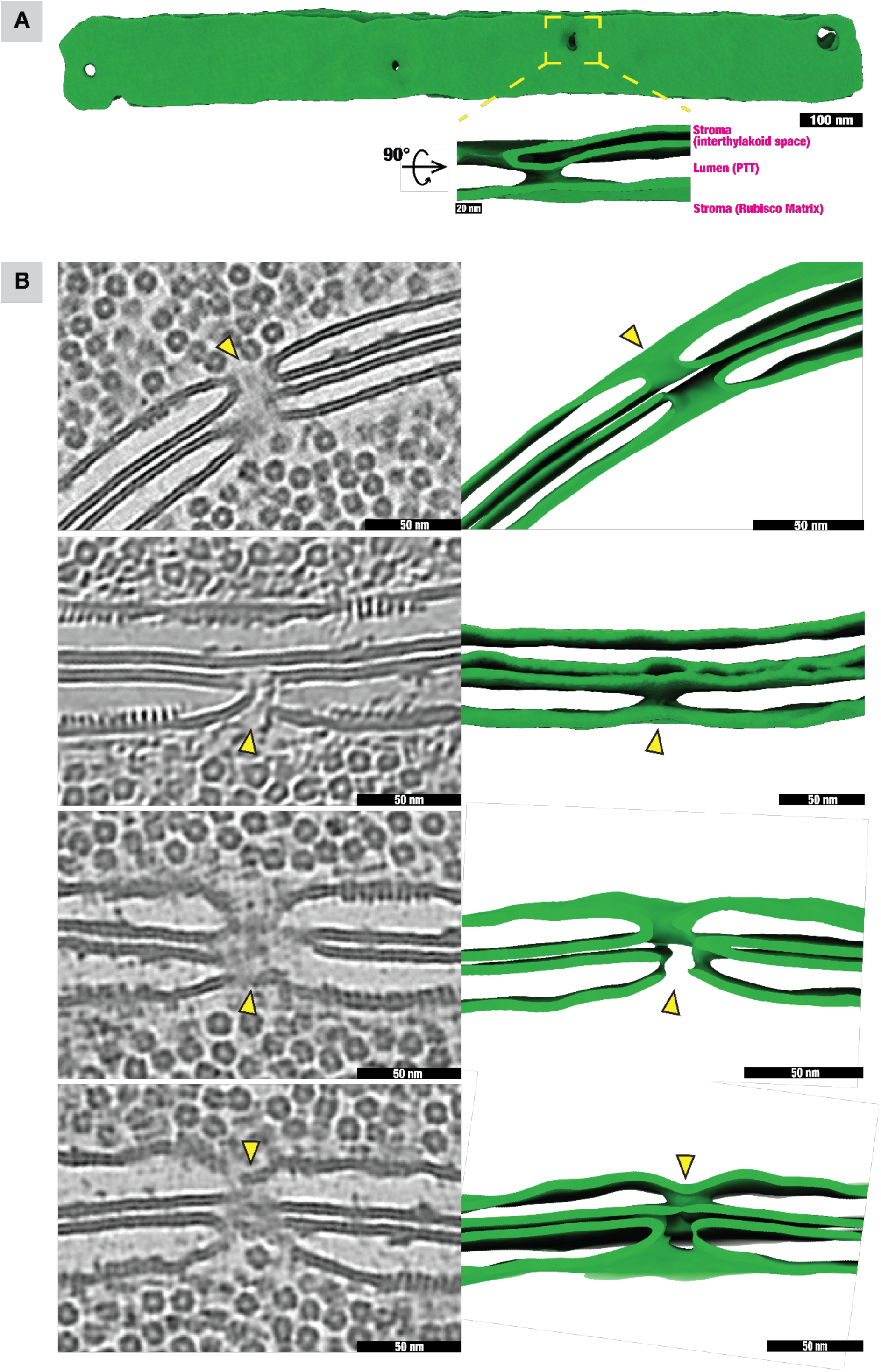
Cryo-ET reveals unique fenestrations in pyrenoid-traversing thylakoid (PTT) membranes of *Chlorella*. a) Membrane segmentation shows multiple fenestrations along PTTs. These holes form conduits that connect the interthylakoid stromal space to the pyrenoid matrix. b) Representative denoised tomographic slices (left) and corresponding membrane segmentations (right). Yellow arrowheads indicate fenestrations forming apparent stroma-matrix connections. Labels: S, interthylakoid stroma; L, PTT lumen; M, pyrenoid matrix.

**Extended Data Fig. 5.**
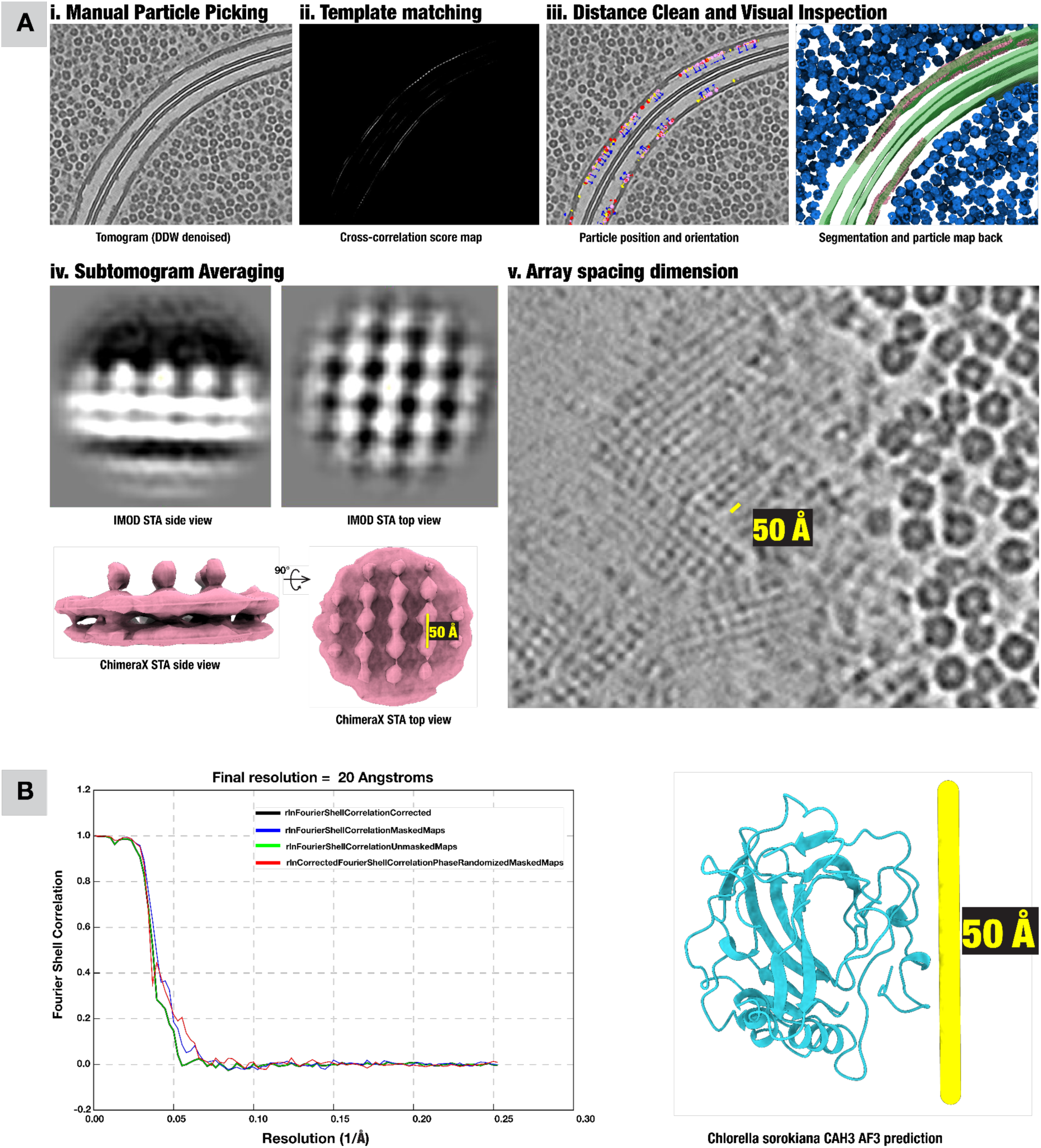
Subtomogram averaging workflow of the putative CAH3 pyrenoid membrane square lattice array. A i) Representative denoised tomographic slice used for initial manual particle picking. ii) Template matching output from initial average shown as a cross-correlation score map. iii) Candidate particle positions and orientations after distance cleaning and visual inspection. Lattice particles and Rubisco mapped back together with the membrane segmentation. iv) Subtomogram average shown from side and top views (top row: slices through map, bottom row: isosurface), highlighting a ∼50 Å lattice repeat. v) Tomographic slice perpendicular to the membrane surface, showing the square array from a top view. B) Fourier shell correlation indicating a final resolution of 20 Å. AlphaFold3 prediction of CsCAH3 showing a similar size to one subunit of the array.

**Extended Data Fig. 6.**
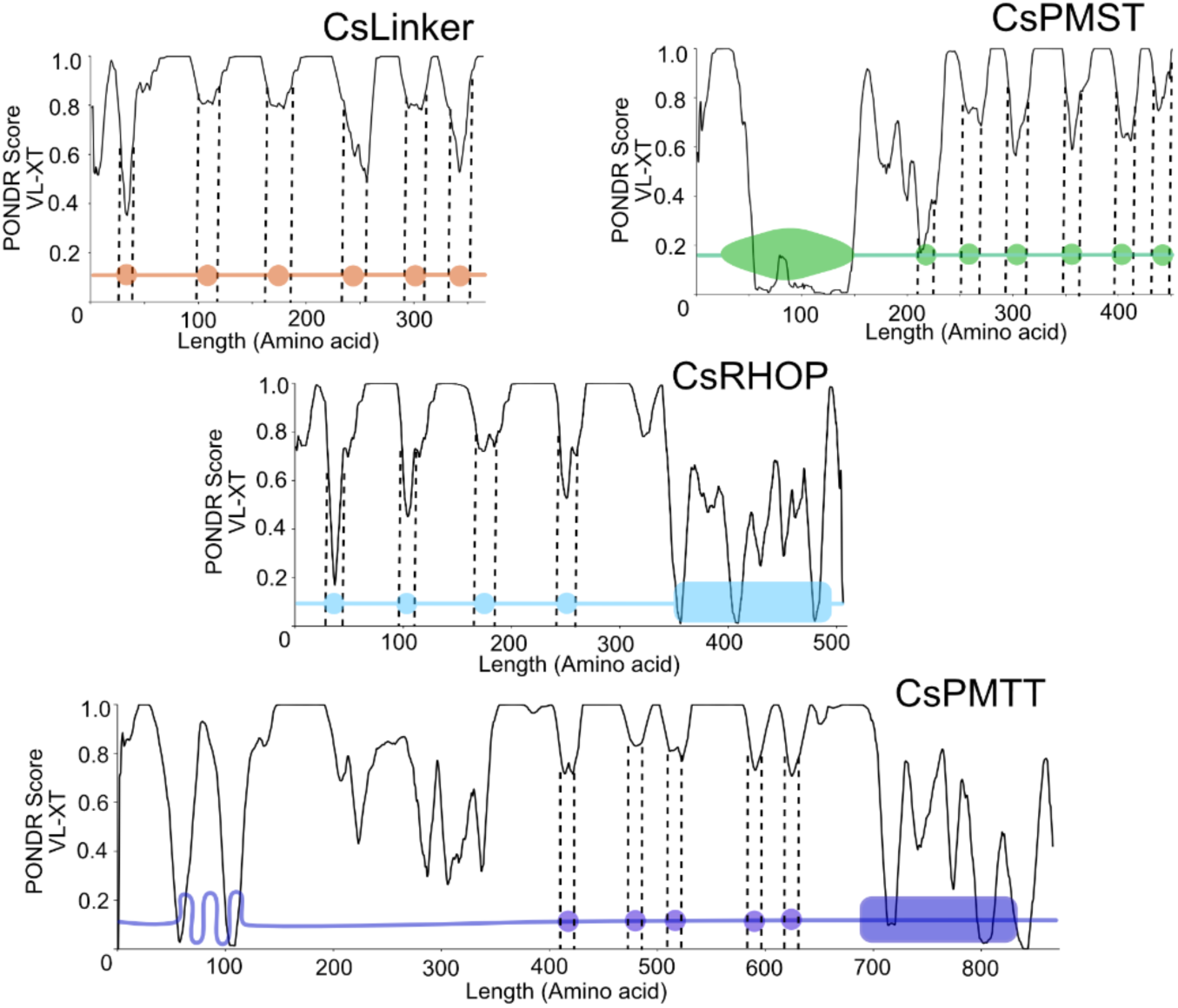
*Chlorella* pyrenoid proteins that contain RBMs that were identified through BLAST search of the CsLinker RBM against the *Chlorella* proteome (CsRHOP and CsPMTT) or through BLAST of search of the CBM20 domain of SAGA1 (CsPMST). PONDR score for predicted disorder of the proteins is shown on the y-axis.

**Extended Data Fig. 7:**
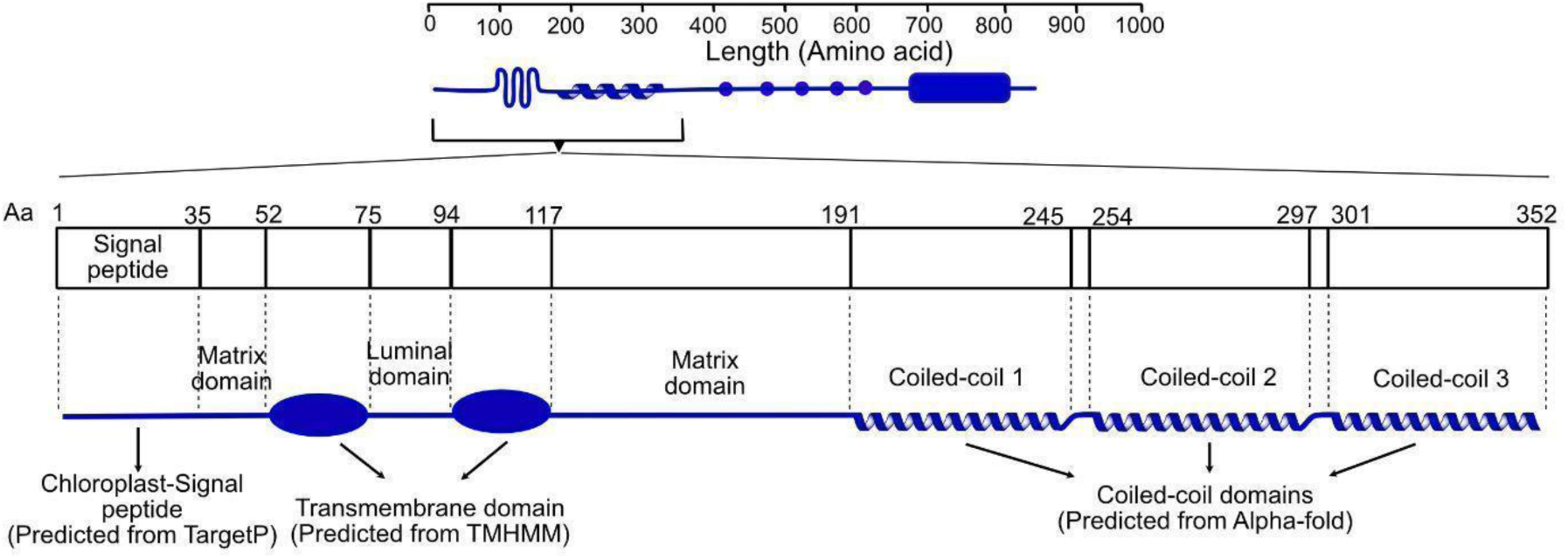
Secondary structure of PMTT with the N-terminal region shown in detail. A chloroplast signal peptide (amino acids 1–35) is predicted by TargetP. Transmembrane topology was predicted using TMHMM on the first 200 amino acids, revealing a short matrix-exposed region (35–52), a first transmembrane domain (52–75), a luminal loop (75–94), and a second transmembrane domain (94–117) extending the remainder of the protein into the matrix. Three coiled-coil domains (191–352) are predicted by AlphaFold.

**Extended Data Fig. 8.**
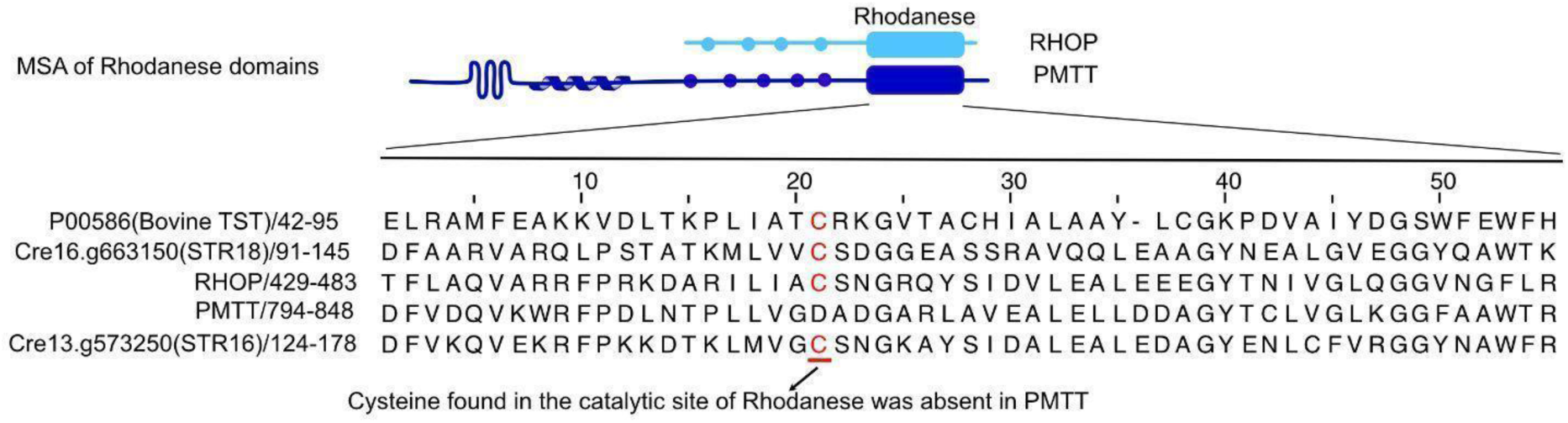
Multiple sequence alignment (MSA) of rhodanese domains from CsPMTT, CsRHOP, and various known active rhodanese-domain proteins shows that CsRHOP contains a conserved cysteine residue within the active site, whereas CsPMTT lacks this cysteine, suggesting its rhodanese domain is likely inactive.

**Extended Data Fig. 9:**
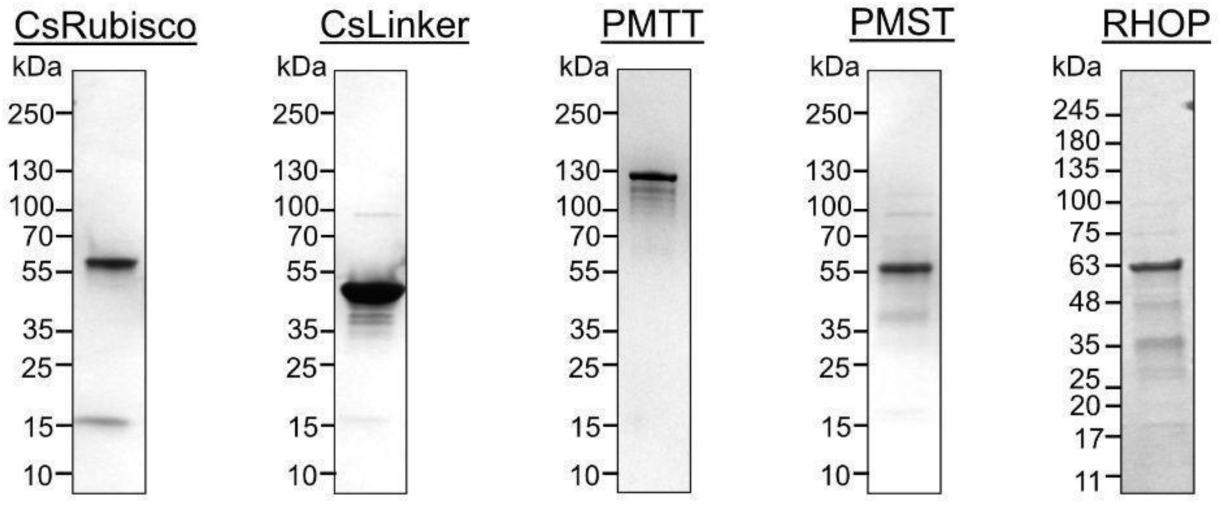
SDS-PAGE of all the purified pyrenoid protein. Approximately 2 μg of each purified protein was resolved by SDS-PAGE to assess purity following the respective purification steps

**Extended Data Fig. 10:**
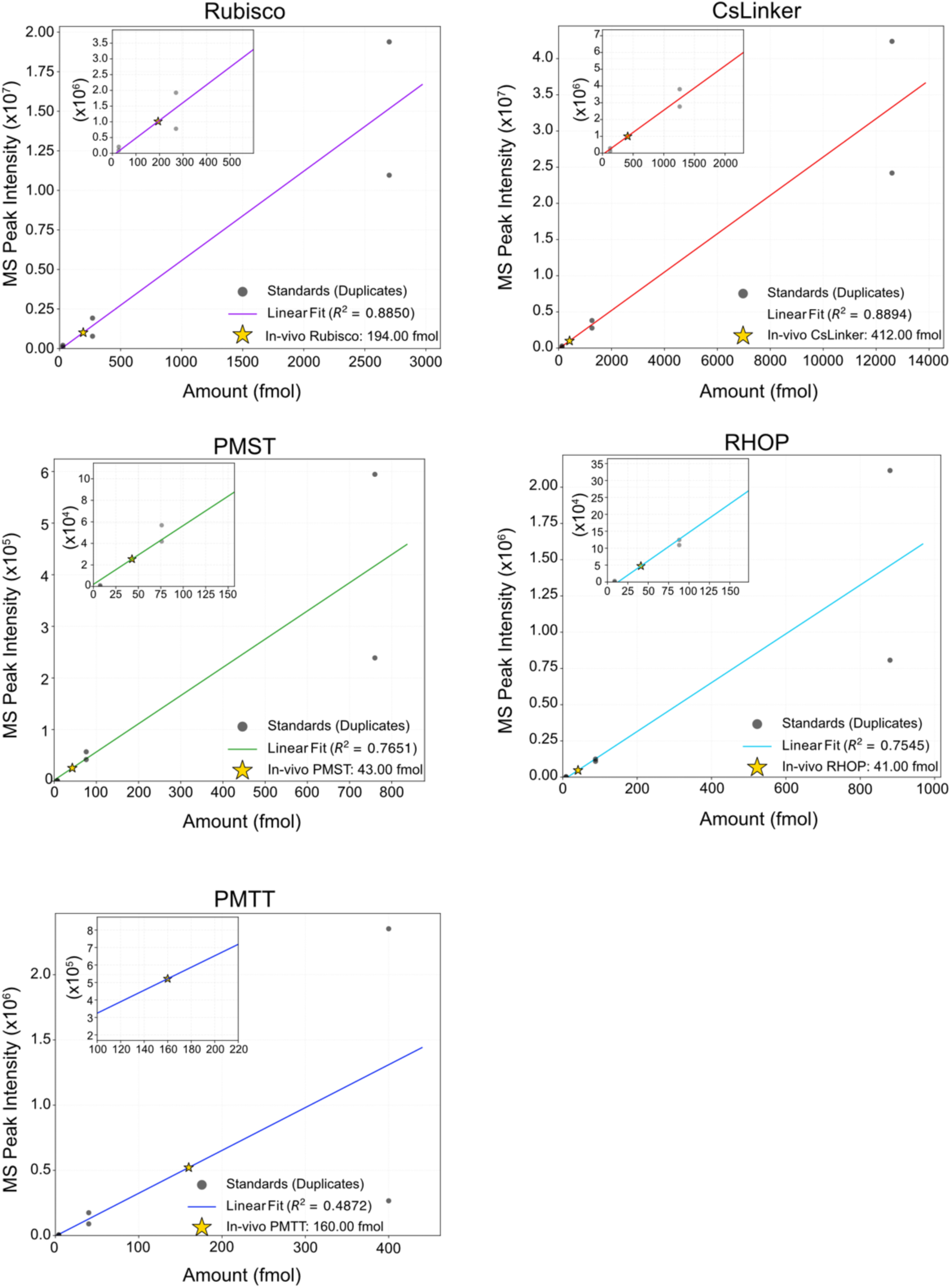
Absolute quantification of CsRubisco, CsLinker, PMST, RHOP and PMTT. a, Standard curves for each protein were generated using purified protein standards to quantify in vivo protein abundance (Star). For purified protein standards, data points represent the S.D. of two technical replicate injections. Rubisco and CsLinker data are obtained from Barrett et al. (2024).

**Extended Data Fig. 11.**
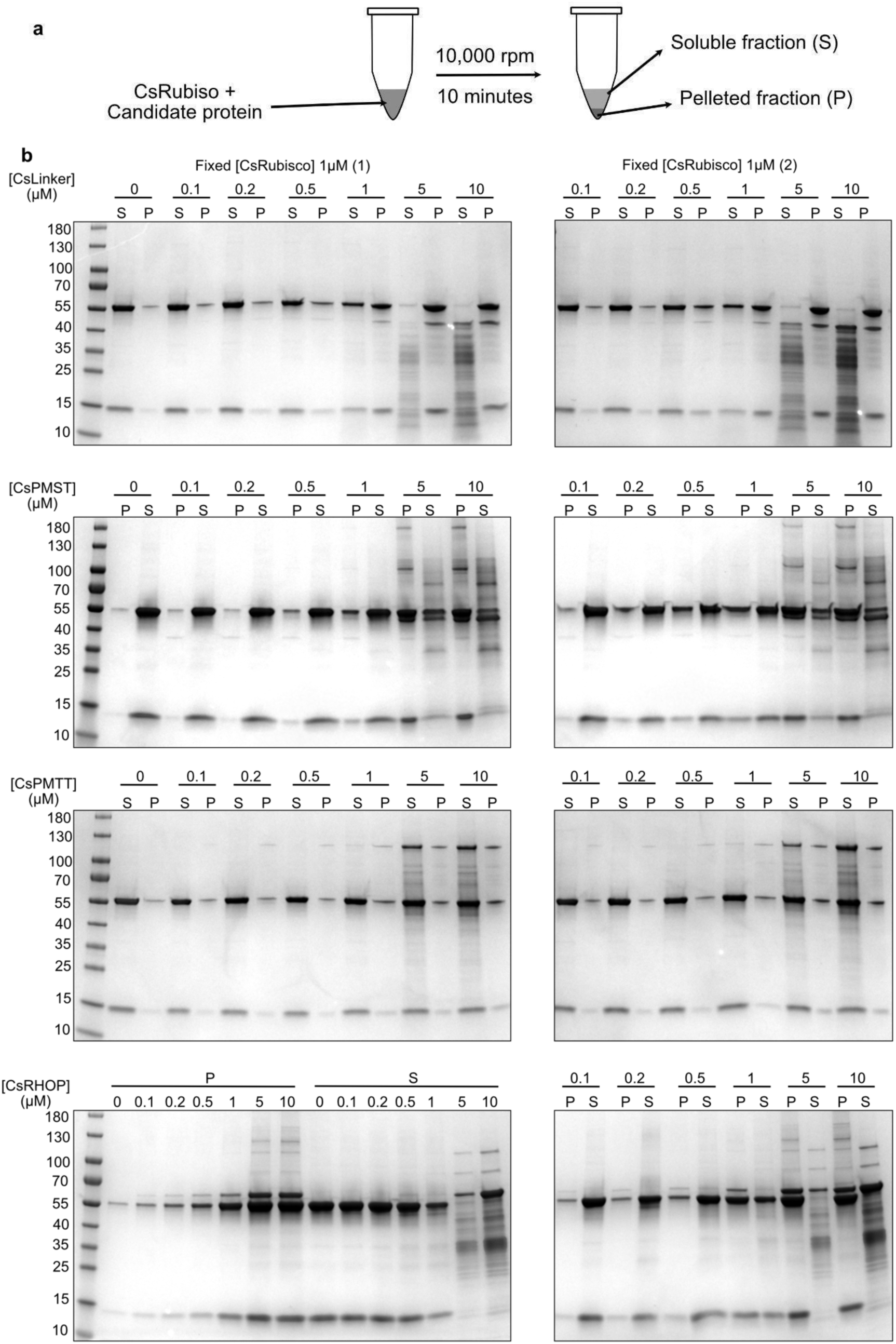
Rubisco sedimentation experiment. a) Schematic of the Rubisco sedimentation assay illustrating the experimental design used to assess the binding and condensation of Rubisco by each candidate *Chlorella* pyrenoid protein. b) SDS-PAGE images of the soluble (S) and pellet (P) fractions obtained for each protein and condition tested. CsRubisco concentration was held constant at 1 μM, and the respective candidate protein concentration for each condition is indicated above each gel. The quantification data presented in Figure 4c are derived from these images.

**Extended Data Fig. 12.**
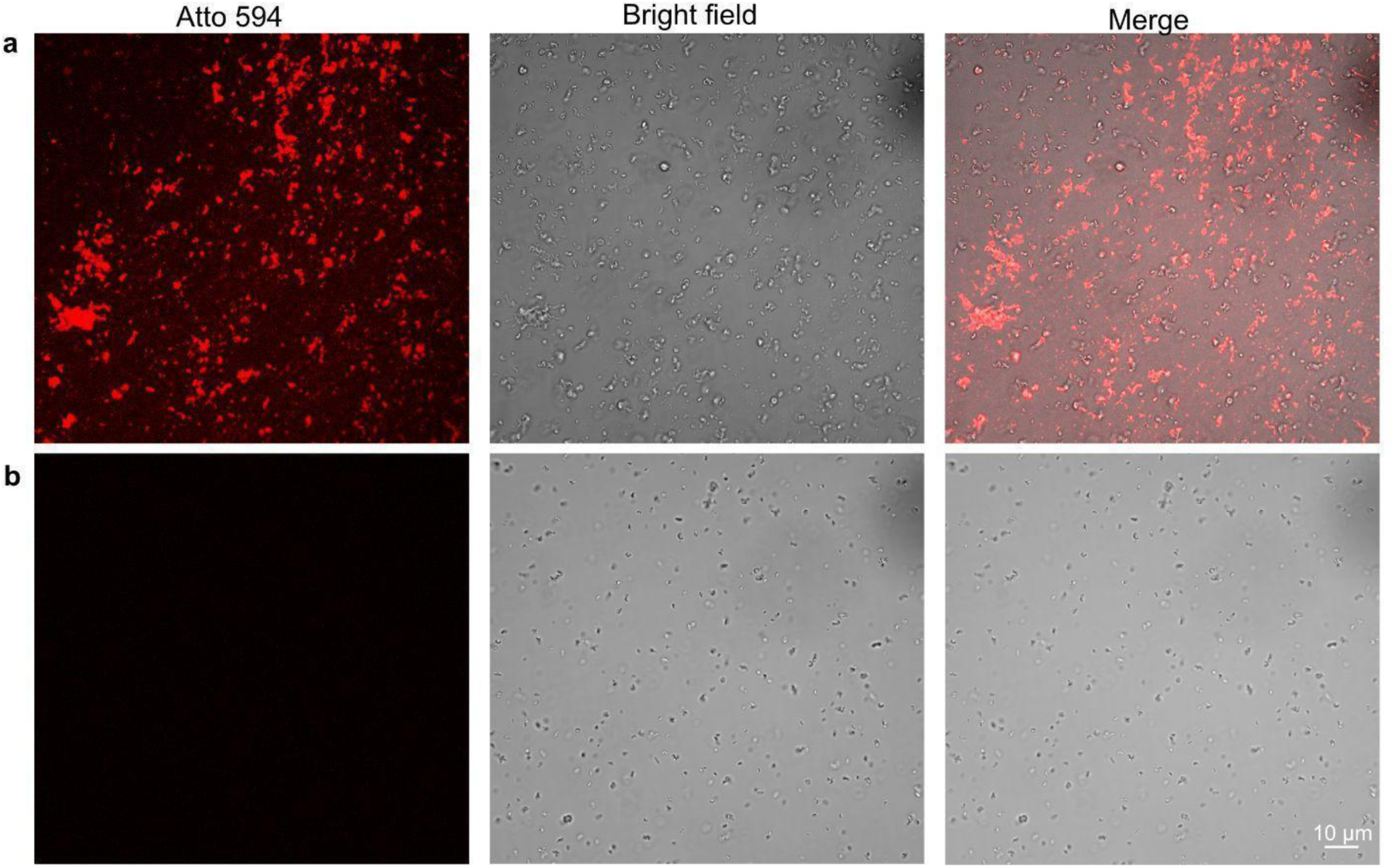
a) Wider confocal image of PMST-Starch and b) ΔCBM20 PMST-Starch interaction. These images were used for quantifying colocalization.

**Extended Data Fig. 13:**
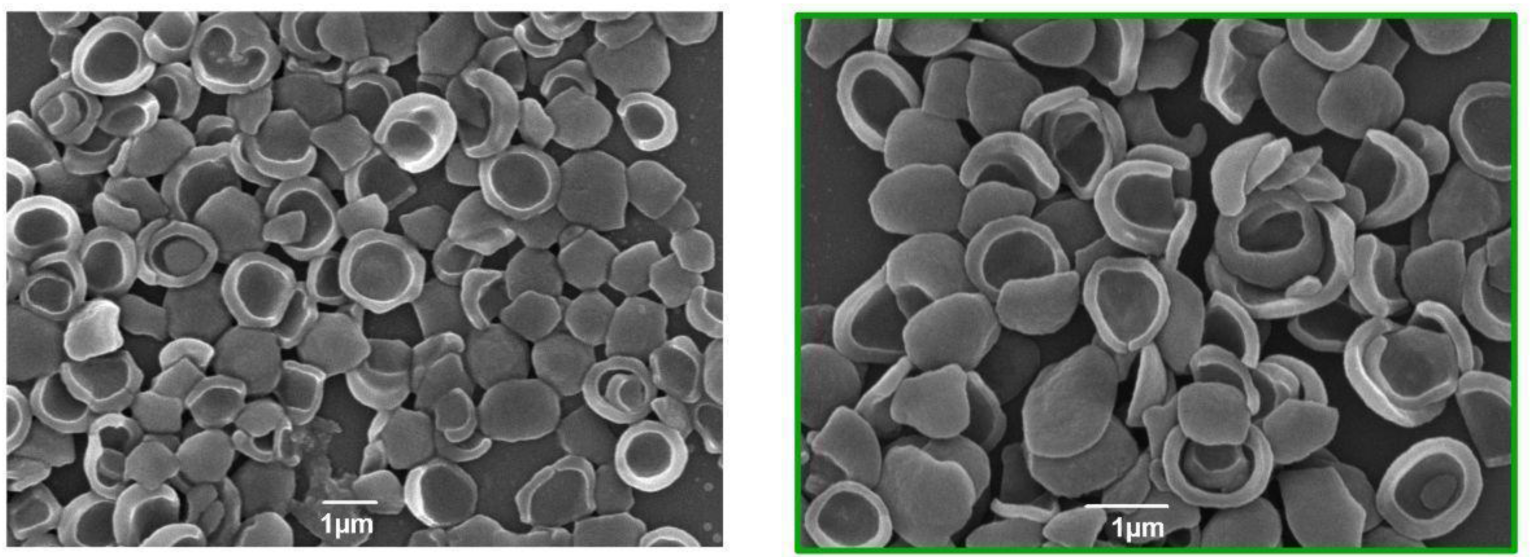
Scanning electron microscopy (SEM) of starch plates isolated from *Chlorella* (left) and starch plates incubated with purified PMST (right). No noticeable structural differences were observed upon PMST treatment.

**Extended Data Fig. 14.**
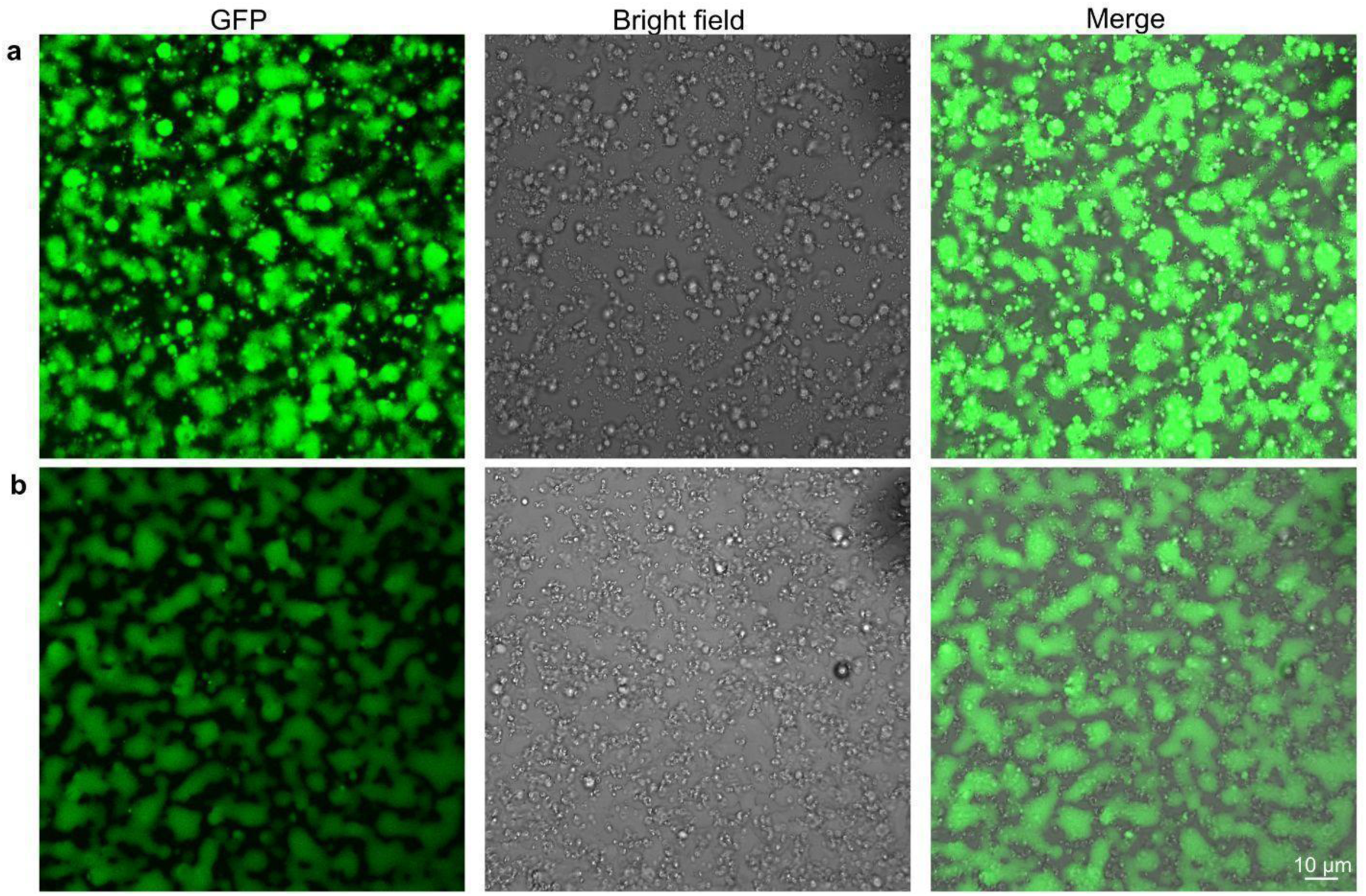
a) Wider confocal image of PMST containing condensate-Starch and b) PMST less condensate-Starch interaction. These images were used for quantifying the fluorescence intensity and colocalisation estimation. Starch plates can be seen in the brightfield as small saucer like structures and condensates are observed as green from the GFP tagged on the Linker protein.

**Extended Data Fig. 15:**
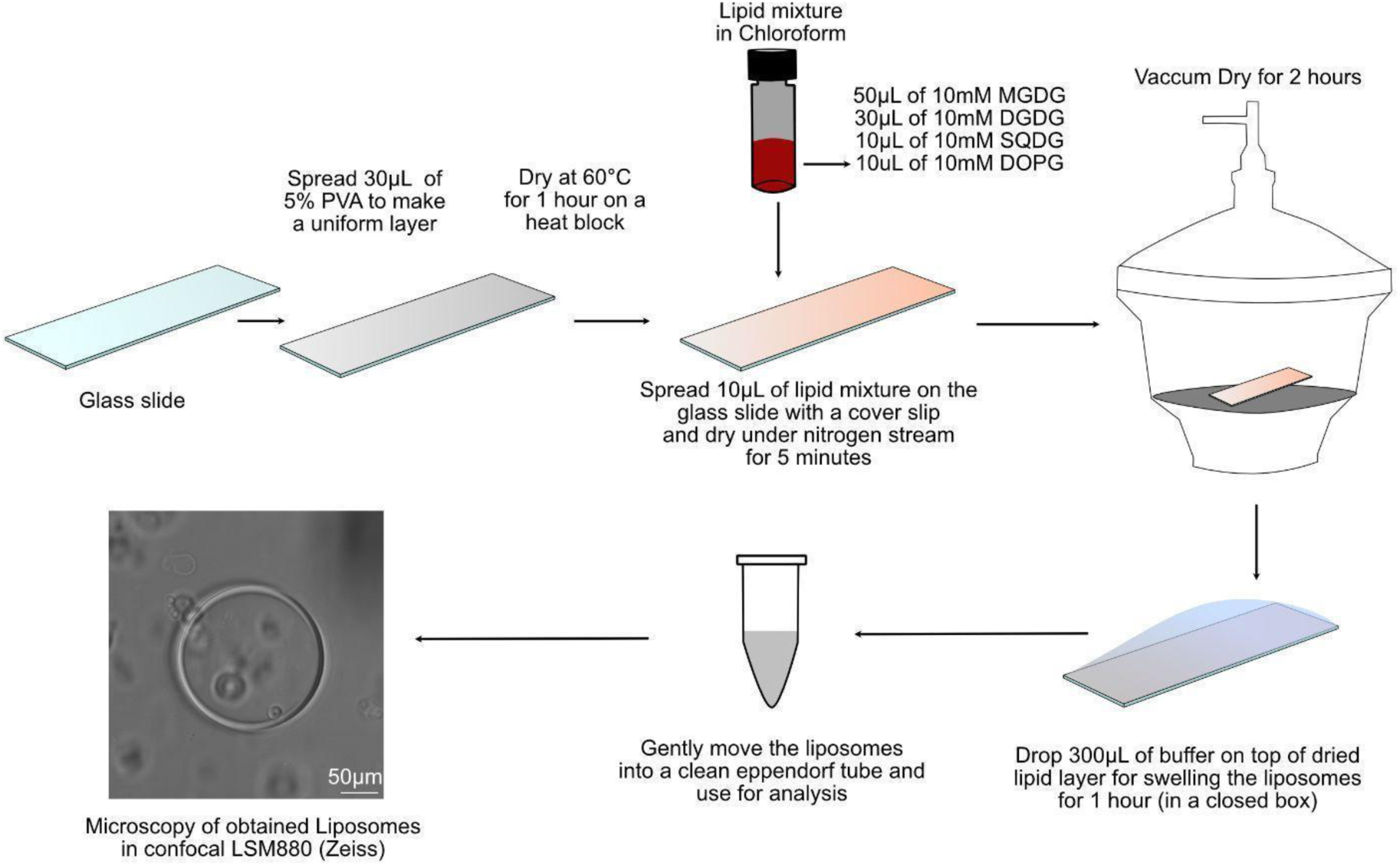
Schematic workflow of developing GUV liposomes with native membrane lipid composition.

**Extended Data Fig. 16:**
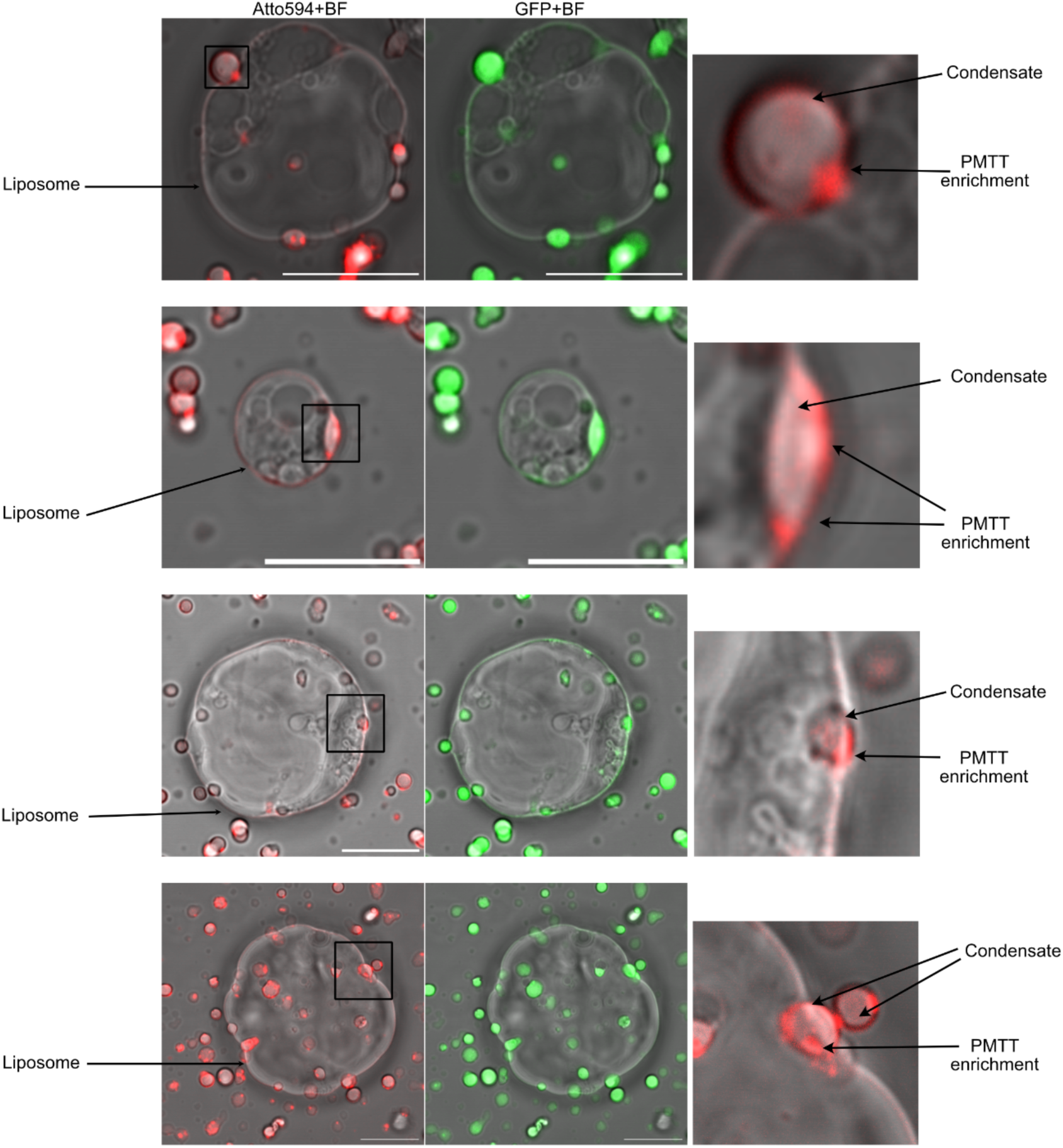
Example images of *native* liposomes mixed with Rubisco-CsLinker condensates and PMTT. Red, PMTT Atto594 (100% labelled PMTT); Green, CsLinker (5%CsLinker-GFP). The bright field (BF) channel shows liposomes and condensates (Scale bar is 10μm).

**Extended Data Fig. 17.**
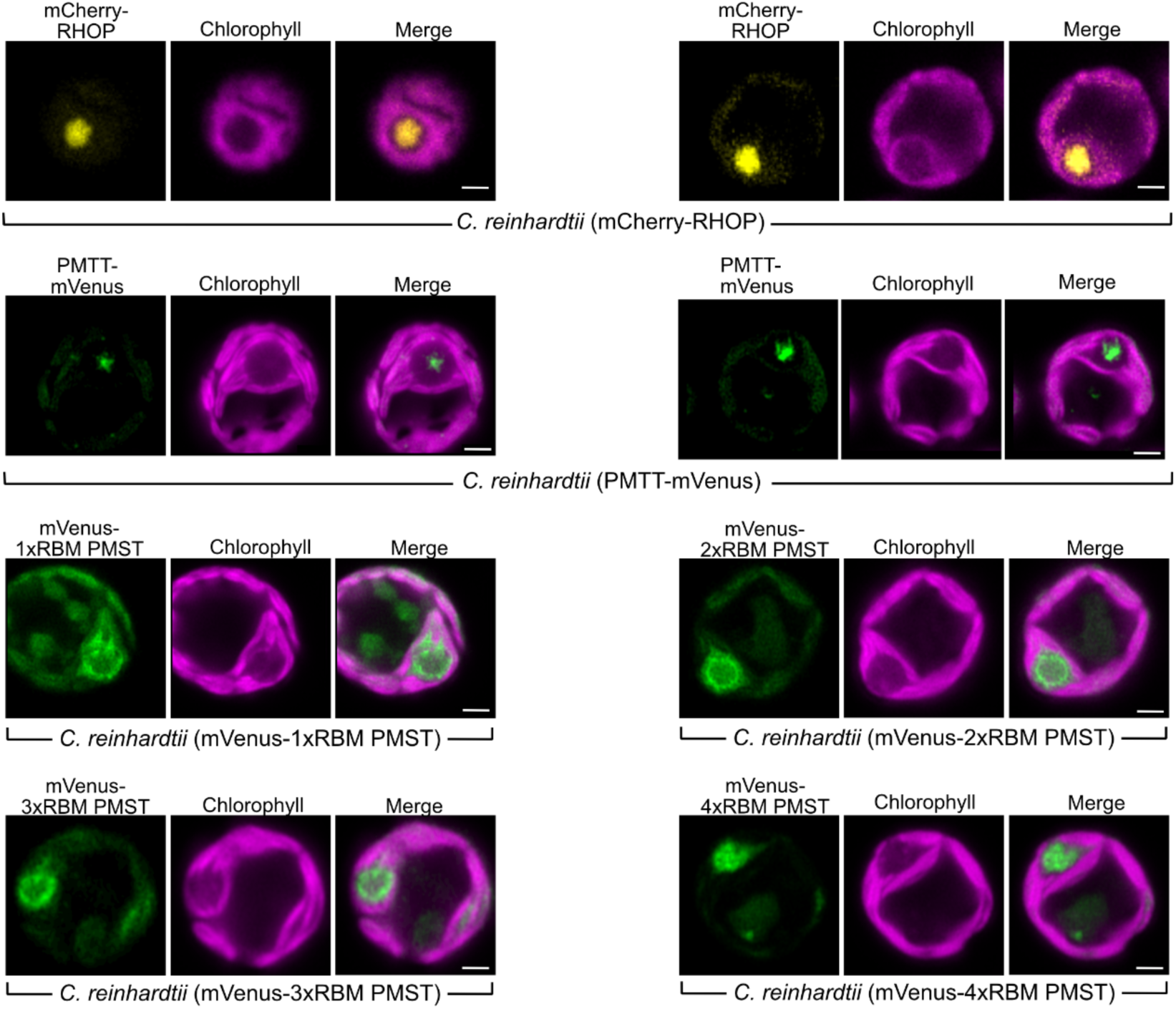
Additional localization images of *Chlorella* pyrenoid proteins expressed from the *Chlamydomonas* chloroplast genome. Scale bar, 2 μm.

**Extended Data Fig. 18.**
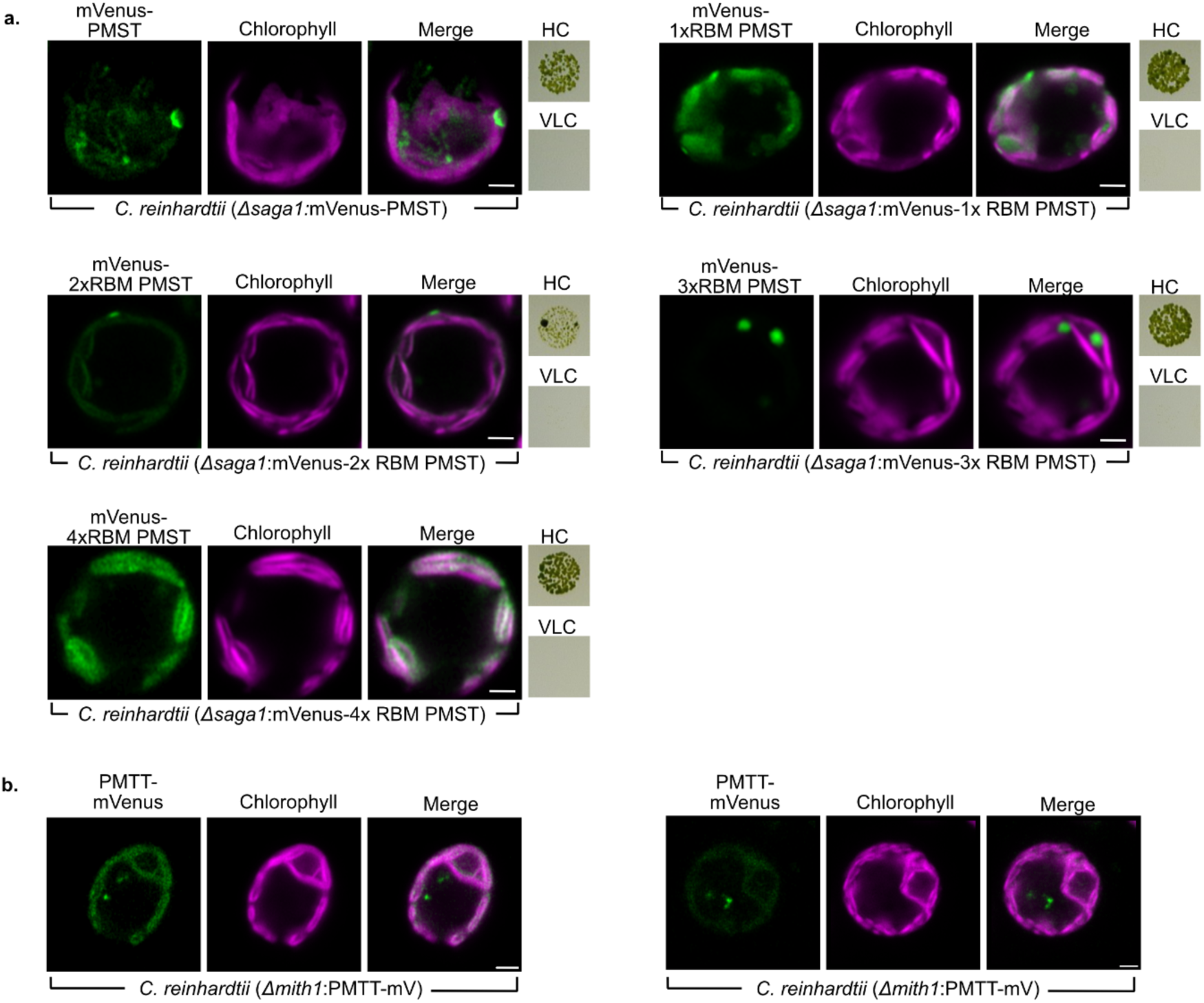
a) Complementation assays of the *saga1* knockout in *Chlamydomonas* with various truncated versions of CsPMST. b) Microscopy of *mith1* expressed with PMTT-mVenus. Scale bar, 2 μm.

**Extended Data Fig. 19.**
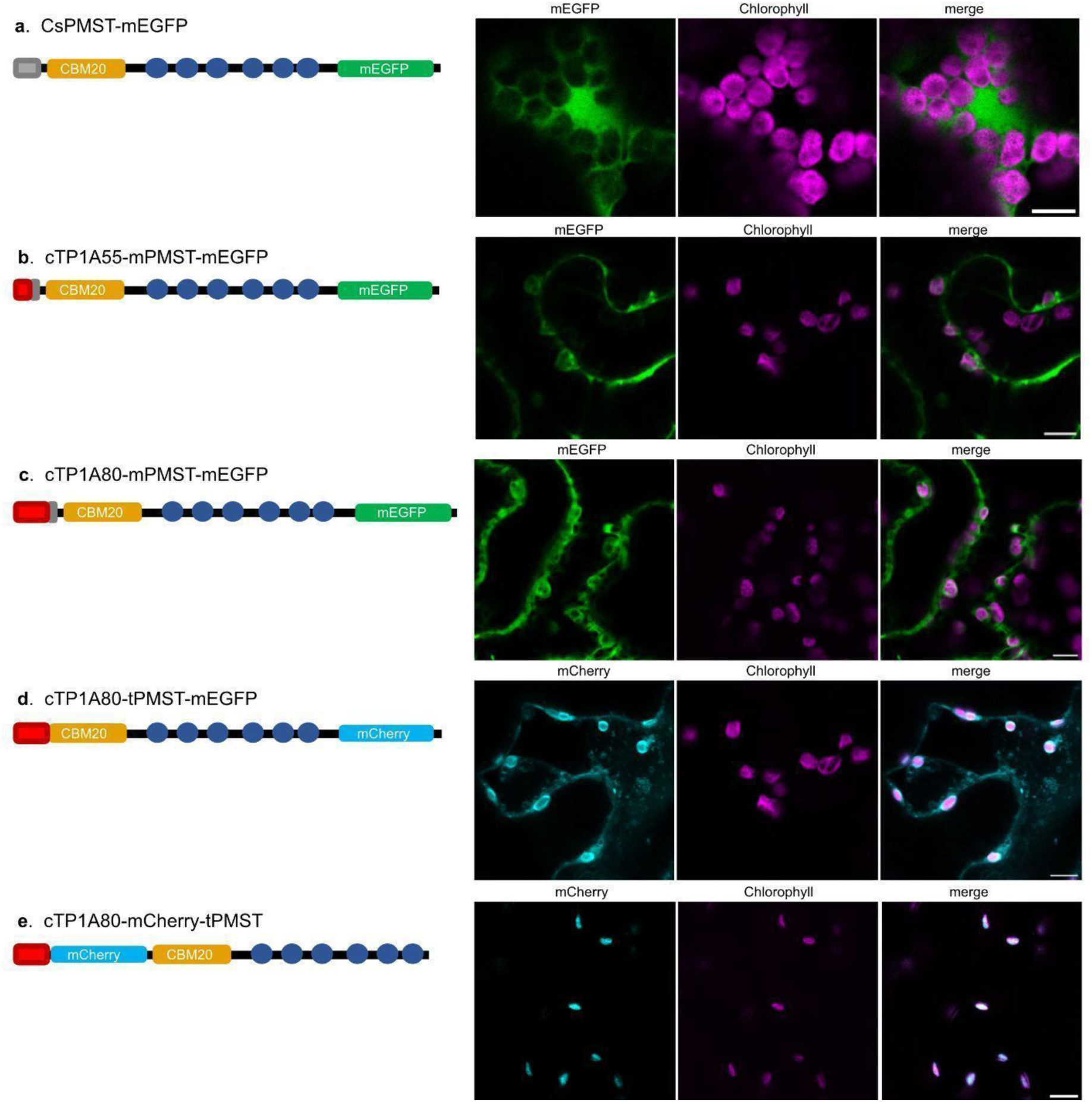
Achieving chloroplastic localization of CsPMST in *N. benthamiana*. CsPMST was codon optimised for expression in plants and expressed in WT *N. benthamiana* plants through infiltration with *Agrobacterium tumefaciens* strain GV3101. Five different constructs were tested through subsequent infiltration experiments to achieve chloroplastic localization. Constructs are represented pictographically where gray/red boxes depict transit peptides, and blue circles represent Rubisco binding motifs. Carbohydrate-binding module 20 (CBM20) and fluorophores are labelled. Scale bar, 10 µm. a) Full-length CsPMST carrying its native chloroplast transit peptide (cTP, grey) fused to mEGFP. b) Mature PMST (mPMST) lacking its native cTP (42 residues removed) and fused to the Arabidopsis RbcS1A cTP (cTP1A55, red). c) A longer version of the Arabidopsis RbcS1A cTP (cTP1A80) was fused to mPMST. d) PMST truncated up to 50 residues from the start codon (tPMST) and fused to cTP1A80. The mEGFP fluorophore was changed to mCherry to allow for visualization of starch with fluorescein staining. e) cTP1A80 and mCherry fused to N-terminal end of tPMST, with a linker sequence (9 residues) added between mCherry and tPMST.

